# The Function of *Efhd1*^+^ Telocytes in the Synovial Lymphatic System and Inflammatory-Erosive Arthritis

**DOI:** 10.1101/2025.11.03.685859

**Authors:** Yue Peng, H. Mark Kenney, Andriy Kobryn, Sean Lydon, Karen L. de Mesy Bentley, Lianping Xing, Benjamin D. Korman, Christopher T. Ritchlin, Edward M. Schwarz

**Affiliations:** Center for Musculoskeletal Research, University of Rochester Medical Center; Department of Pathology & Laboratory Medicine, University of Rochester Medical Center; Department of Medicine, Division of Allergy, Immunology, Rheumatology, University of Rochester Medical Center

**Keywords:** Telocyte, Mast Cell, Lymphatic Vessel, Electron Microscopy, Inflammatory-Erosive Arthritis, Animal Models

## Abstract

Resting collecting lymphatic vessels (cLVs) sense edema in distal joints and initiate contractions via unknown mechanisms. Rheumatoid arthritis (RA) patients have lymphatic drainage deficiencies from affected joints, and defects in the synovial lymphatic system exacerbate inflammatory-erosive arthritis in animal models. To understand this, we generated *Efhd1*-CreER^T2^ and *Myoc*-CreER^T2^ mice for cell-specific genetic gain and loss of function studies. These mice were crossed with tdTomato reporter (Ai9) mice, and studies showed selective tamoxifen-induced transgene expression in CD31^−^/CD34^+^ telocyte-like cells in knee and ankle synovium, and in networks physically associated with mast cells proximal to popliteal lymphatic vessels (PLVs). Consistent with the known loss of CD31^−^/CD34^+^ telocyte in RA synovium, these cells were also decreased in TNF-tg knees and partially recovered by anti-TNF treatment. Ultrastructural and gene expression studies confirmed a distinct telocyte phenotype versus closely related fibroblasts. *In vivo* depletion studies in tamoxifen-treated *Efhd1*-CreER^T2^ and *Myoc*-CreER^T2^ mice crossed to diphtheria toxin alpha-floxed (DTA^flox^) mice demonstrated telocyte requirements for physiologic lymphatic drainage and resolution of joint inflammation and focal erosions from zymosan-induced arthritis in the knee. *In vitro* studies demonstrated increased sensitivity to osmotic shock and decreased motility versus fibroblasts, and telocyte potential to differentiate into myofibroblasts on stiff matrix. Collectively, these findings support a model of joint homeostasis in which osmotic pressure-sensing telocyte networks extend from the synovium into mast cells proximal to joint-draining cLVs, and telocyte loss is associated with defects in the synovial lymphatic system and increased susceptibility to joint inflammation and structural damage from arthritis.

**Teaser:** This study discovered a unique cell type (telocytes) responsible for draining inflammation from arthritic joints.

## INTRODUCTION

The lymphatic system reabsorbs excess interstitial fluid and protein for return to the central veins, and dysfunction is associated with lymphedema in affected tissues, exacerbation of chronic inflammatory diseases, and fibrosis (*1*). While some lymph migrates passively through unidirectional bicuspid lymphatic vessel valves secondary to interstitial fluid pressure and passive compression by adjacent tissue movement (*2*), the majority of lymph is transported by the contractile activity of collecting lymphatic vessels (cLVs) (*3*). Central to cLV functions are tightly regulated contraction frequency changes (*4*), known as ‘pressure-induced lymphatic chronotropy’ (*5*), which range from ∼3 to 20 contractions/min as intraluminal pressure increases from 0.5 to 10 cmH_2_O (*6, 7*) to enable coupled lymph transport with absorption via lymphatic capillaries (*8*). Recently, major advances have greatly enhanced our understanding of intraluminal pressure-induced lymphatic chronotropy and intrinsic cLV pacemaking by lymphatic muscle cells (LMCs) (*5, 6, 9, 10*). Interestingly, LMCs contain functional and molecular characteristics of both myocardium (cardiac muscle) and blood vessels (vascular smooth muscle), as evidenced by concomitant rapid cardiac-like phasic contractions to generate flow and smooth-muscle-like slow, long-lasting tonic contractions to regulate flow (*11*). However, several critical knowledge gaps remain including the identity of cells that sense acute edema in tissue distal to cLVs to trigger enhanced lymphatic clearance by initiating pacemaking signals in LMCs, and the mechanisms that underlie the loss of cLV contractility in lymphedema (*12*).

We previously demonstrated that in rheumatoid arthritis (RA), lymphatic dysfunction is a critical factor in joint inflammation, damage, and flare which can progress independent of autoimmunity (*13*). Evidence of synovial lymphatic system dysfunction in osteoarthritis (OA) also exists (*14*). Studies on innate immune mechanisms identified increased expression of the lymphangiogenic factor vascular endothelial growth factor C (VEGF-C) in monocytes and osteoclasts, together with lymphangiogenesis in inflamed joints (*15*). Subsequent functional studies demonstrated that loss of VEGF-C signaling together with decreased lymphangiogenesis exacerbates inflammatory-erosive arthritis (*16*), while VEGF-C therapy ameliorates inflammatory arthritis by increasing lymphangiogenesis (*17*). Longitudinal MRI (*18, 19*), power Doppler ultrasound (PDUS) (*20*), and near-infrared (NIR) imaging of injected indocyanine green (ICG) dye (*7, 21*), identified three distinct mechanisms of lymphatic dysfunction in murine models of RA. The first is mediated by release of local vasodilators (e.g. nitric oxide) from inflammatory cells that contribute to reduced lymphatic flow (*13, 22*). Another involves accumulation and translocation of B cells into the sinuses of joint-draining lymph nodes, obstructing passive lymphatic flow (*19, 21, 23*). Finally, we observed the loss of cLV contractions during arthritic progression in the setting of chronic joint inflammation (*20*). In support of the translational relevance of these findings are data demonstrating that RA patients display similar lymphatic dysfunction, including reduced cLV contractility and lymphatic clearance compared to healthy controls (*24–26*), which is now a high priority area of research.

Mast cells are recognized as effective regulators of LMC contractility due to their anatomical proximity and ability to produce, store, and release various inflammatory and vasoactive mediators (*27*). Additionally, mast cells are positioned throughout the RA synovial sublining, where they comprise 5% or more of the expanded synovial cell population, and microanatomic clusters of mast cells are present in pannus tissue proximal to cartilage and bone erosion (*28*). Based on these findings and evidence from preclinical and clinical studies supporting pro-inflammatory and catabolic roles for mast cells in RA (*28, 29*), we investigated mast cell effects on joint-draining lymphatics and inflammatory-erosive arthritis in tumor necrosis factor-transgenic (TNF-tg) mice (*30*) and their wild-type (WT) littermates. We found that popliteal lymphatic vessels (PLVs) are surrounded by an expanded frequency of MCT^+^/MCPT1^+^/MCPT4^+^ mast cells in TNF-tg mice, and the percentage of peri-PLV degranulating mast cells was inversely correlated with lymphatic clearance (*31*). We also identified a unique population of MCT^+^/MCPT1^−^/MCPT4^−^ mast cells embedded within the PLV cellular architecture, but the function of these cells remains unknown (*31*). Genetic and pharmacologic mast cell loss of function experiments in TNF-tg mice demonstrated severe lymphatic defects, which were associated with exacerbated inflammatory-erosive arthritis (*31*). Collectively, these findings led us to surmise that while MCT^+^/MCPT1^+^/MCPT4^+^ mast cells have a well-established role in inflammation, particular mast cell subtypes are also required for normal lymphatic function, proposed to be mediated by MCT^+^/MCPT1^−^/MCPT4^−^ mast cells embedded within PLVs. However, several unanswered questions remain including: i) the anatomy and physiology of mast cell integration into the synovial lymphatic system, ii) the mechanisms that facilitate recognition of increased joint fluid by the distal cLV, iii) the resulting increased contraction frequency during the early phase of inflammation, and iv) how this compensatory response is lost during chronic arthritis, remain be elucidated.

To address these questions, we developed conditional-inducible transgenic mice for genetic gain and loss of function studies initially considered to specifically target LMCs. Single-cell RNA sequencing (scRNAseq) studies identified three genes (*Efhd1, Myoc & Pla1a*) in stromal cell population selectively expressed in PLVs compared to adjacent blood vessels (*32*). We then generated three transgenic mouse lines in which a *CreER^T2^* cassette was knocked into these genes, and the resulting animals were crossed to B6.Cg-*Gt(ROSA)26Sor^tm9(CAG-tdTomato)Hze^*/J (Ai9) (*33*) and B6.129P2-*Gt(ROSA)26Sor^tm1(DTA)Lky^*/J (DTA) (*34*) mice for lineage tracing and loss of function studies, respectively. Herein, we describe the construction of these models, and the serendipitous finding of tamoxifen-inducible gene expression in peri-PLV spindle-shaped cells with very long cytoplasmic extensions that were physically integrated into mast cells, while there was no expression in the predicted LMC population (*32*), which has been further described in more recent scRNAseq studies (*9*). We hypothesized the identified peri-PLV spindle-shaped cells to be telocytes, which are unique mesenchymal cells found in most tissues and are characterized by a small cell body and long, thin, cytoplasmic extensions called telopodes (*35*). Of note are the various telocyte functions that include extracellular matrix (ECM) synthesis for tissue structural support, cell-to-cell communication, regulation of immune responses, and pacemaking of smooth muscle contractions (*36*). To confirm these characteristics supporting telocyte identity and determine the function of telocytes within the synovial lymphatic system, we performed various *in vivo* and *in vitro* studies of the tdTomato-positive (tdT^+^) cells. Together, we tested the hypotheses that: 1) networks of contiguous telocytes extend from ankle synovium to PLVs, 2) telocytes are functionally distinct from lymphatic fibroblasts, and 3) synovial telocytes differentiate into myofibroblasts when grown on stiff ECM.

## RESULTS

### Generation of Novel CreER^T2^ Mouse Models for Genetic Targeting of Telocytes in the Synovial Lymphatic System

We recently described single-cell RNA sequencing (scRNAseq) analysis of lymphatic vessels in which we identified *Efhd1, Myoc,* and *Pla1a* as PLV-selective genes (*32*). Based on these findings, we generated three knock-in mouse lines in which a *CreER^T2^* cassette was inserted into these genes of embryonic stem cells (Fig. 1a), and the resulting mice were crossed to Ai9 reporter mice to assess tamoxifen-induced in vivo gene targeting (Fig. 1b & c). *Ex vivo* whole-mount immunofluorescent microscopy (WMIFM) revealed the presence of elongated tdT^+^ cells that were preferentially clustered in networks around PLVs vs. blood vessels from tamoxifen-treated *Efhd1*-CreER^+/-^x Ai9^+/-^ and *Myoc*-CreER^+/-^ x Ai9^+/-^ mice (Fig. 1d). Tamoxifen treatment of younger mice (P10-P13) resulted in a similar pattern of PLV adventitial tdT^+^ cells, suggesting that they maintain their identity from early postnatal development (Supplemental Fig. 1a-b). These tdT^+^ cells were not found in tissues from tamoxifen-treated *Pla1a*-CreER^+/-^ x Ai9^+/-^ mice, and these mice were not further analyzed (Supplemental Fig. 1b).

**Fig. 1.**
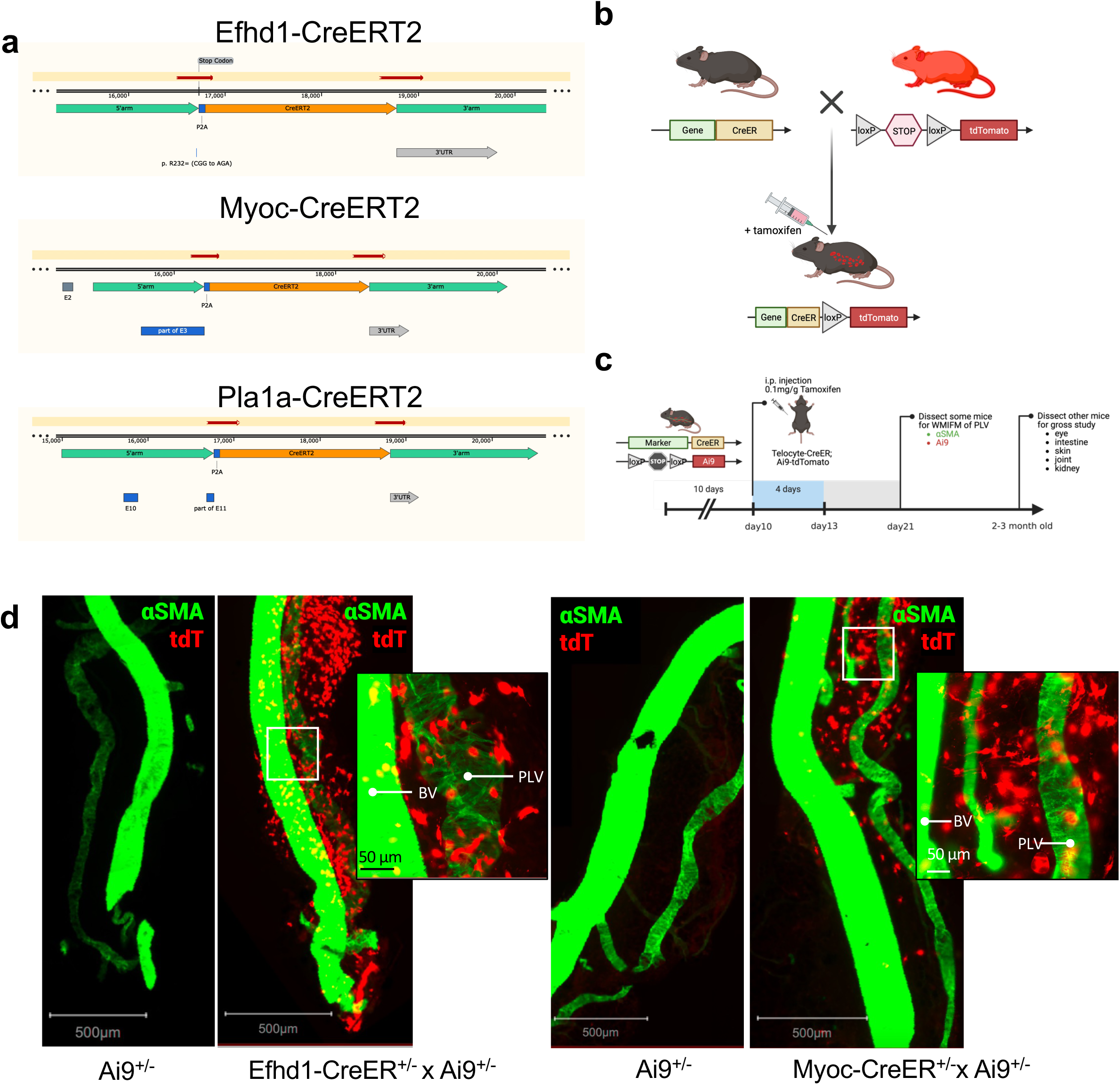
Generation and validation of CreERT2 transgenic mice with tamoxifen-inducible gene expression in PLV adventitial cells. **a** Schematic illustration of the P2A-CreERT2 targeting vectors that were used to generate the knock-in transgenic mice as described in Methods. **b** Schematic illustration of tdT reporter studies performed to assess tissue-specific tamoxifen-inducible gene expression in the CreERT2 knock-in transgenic mice in which founder lines were crossed with B6.Cg-*Gt(ROSA)26Sor^tm9(CAG-tdTomato)Hze^*/J (Ai9) mice to generate *Efhd1*, *Myoc* and *Pla1a* CreERT2^+/-^ x Ai9^+/-^ test mice and single-transgenic Ai9^+/-^littermate controls. **c** An illustration of the tamoxifen dosing regimen (daily intraperitoneal injections on postnatal days 10–13) to the 3 lines of male and female mice (n ≥ 5) is presented with the standard time of sacrifice at postnatal day 21. **d** WMIFM was performed on PLV immunostained with FITC-conjugated anti-αSMA antibodies from the indicated tamoxifen-treated mice. Representative 4x images of tdT^+^ cells (red) surrounding αSMA^+^ (green) in Efhd1CreER^+/-^ x Ai9^+/-^ and MyocCreER^+/-^ x Ai9^+/-^ mice after tamoxifen induction, but not in CreER negative controls.

Interrogation of the PLV WMIFM revealed two distinct tdT^+^ populations of fibroblastic cells that contained very long dendritic processes phenotypically consistent with telopodes (Fig. 2a). One population extended longitudinally along the PLV (Fig. 2b), and the other existed in telocyte arrays within adipose tissue adjacent to PLVs (Fig. 2c). To gain wholistic insight on the extent of tdT^+^ cell arrays along the PLV, we performed WMIFM on tissues from tamoxifen treated *Efhd1*-CreER^T2^ x Ai9 mice with antibodies against αSMA and CD34 to label the vascular smooth muscle and mast cells. Confocal images of soft-tissue extending from the ankle to the knee containing the PLVs were reconstructed in 3D and stitched together (Figs. 2d-h). The high-power images confirmed the presence of tdT^+^ cell arrays adjacent to PLVs that contained CD34^+^ cells with a mast cell morphology. The low-resolution images revealed the networks of tdT^+^ cells that paralleled the vessels from the ankle synovium to cLV.

**Fig. 2.**
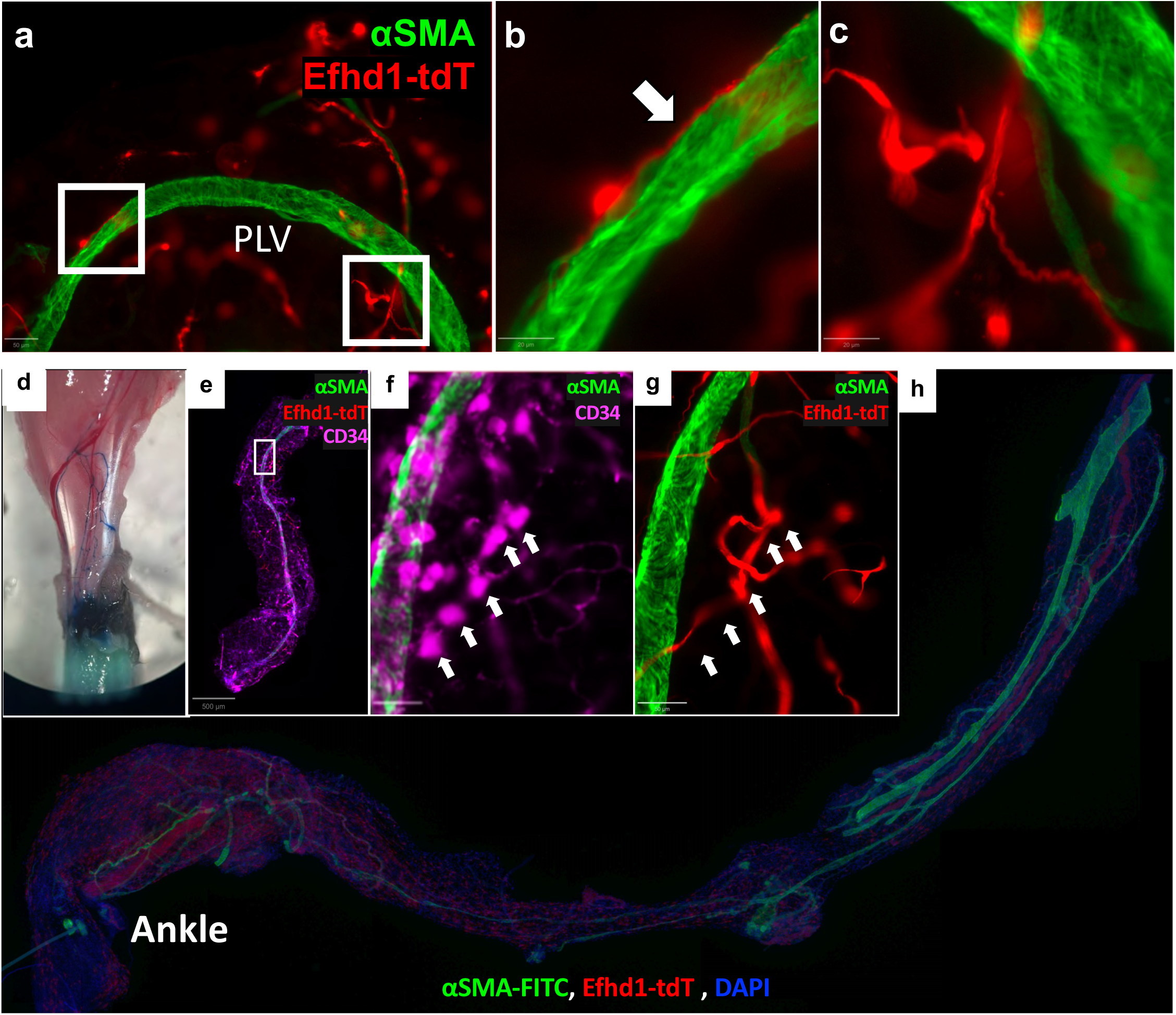
Efhd1-CreERT2 targets two populations of PLV adventitial cells that are adjacent to mast cells. **a** WMIFM of PLV from tamoxifen-treated Efhd1-CreER^+/-^x Ai9^+/-^ mice was performed as described in Figure 1 and a 4x image is show with 10x images of the boxed ROIs highlighting **b** a tdT^+^ telocyte-like cell with its telopode (arrow) aligned along the surface of the PLV, and **c** a network of tdT^+^ telocyte-like cells in the adipose tissue adjacent to the vessels. **d** Evan’s blue dye was injected into the footpad of a tamoxifen-treated *Efhd1*-CreERT2 x Ai9 mouse to image the PLVs for dissection, and the tissue was immunostained with fluorescent antibodies against CD34 and αSMA, and counterstained with DAPI. **e-g** Low power and high-power confocal microscopy images of the ROI (box) confirm immunostaining of the mast cells elaborating the tdT^+^ network along the PLV (green). **h** Eight fluorescent low power images of an immunostained full-length PLV with associated tissues were stitched together to imaging telocyte networks that extends from capillaries in the ankle synovium to cLV near the knee.

Previous investigations of lymphatic dysfunction revealed an enrichment of mast cells around lymphatic vessels (*31*), which prompted us to examine these distinct tdT^+^ cells and their interactions within the peri-PLV tissue more closely. Scrutiny of transmission electron microscopy (TEM) images of PLVs revealed elongated cells in tamoxifen-treated *Efhd1*-CreER^+/-^x Ai9^+/-^ and *Myoc*-CreER^+/-^ x Ai9^+/-^ mice with morphological characteristics consistent with the tdT cells identified by WMIFM (Fig. 3a). Moreover, these cells have characteristic ultrastructural features of telocytes (*35*) (i.e. spindle-shaped polygonal cells with a large single nucleus and scant cytoplasm and thinly attenuated cytoplasmic extensions). These cells were found to form intimate contacts with mast cells in what appears to be specialized junctional structures at their plasma membrane interfaces suggesting active intercellular communication (Fig. 3a). To validate these observations and better characterize the spatial relationships between telocytes and mast cells, we conducted 3D immunofluorescent confocal microscopy. This analysis demonstrated that mast cells form direct contacts with the two tdT^+^ telocyte-like populations (Fig. 3b and Supplemental Video 1). The spatial distribution and interaction patterns observed through 3D confocal imaging precisely matched our WMIFM and TEM observations, confirming that both mural and peri-PLV telocyte populations engage in direct cellular interactions with mast cells. Taken together, these findings suggest a previously unrecognized cellular partnership between telocytes and mast cells along lymphatic vessels, potentially representing a novel signaling axis in cLV regulation.

**Fig. 3.**
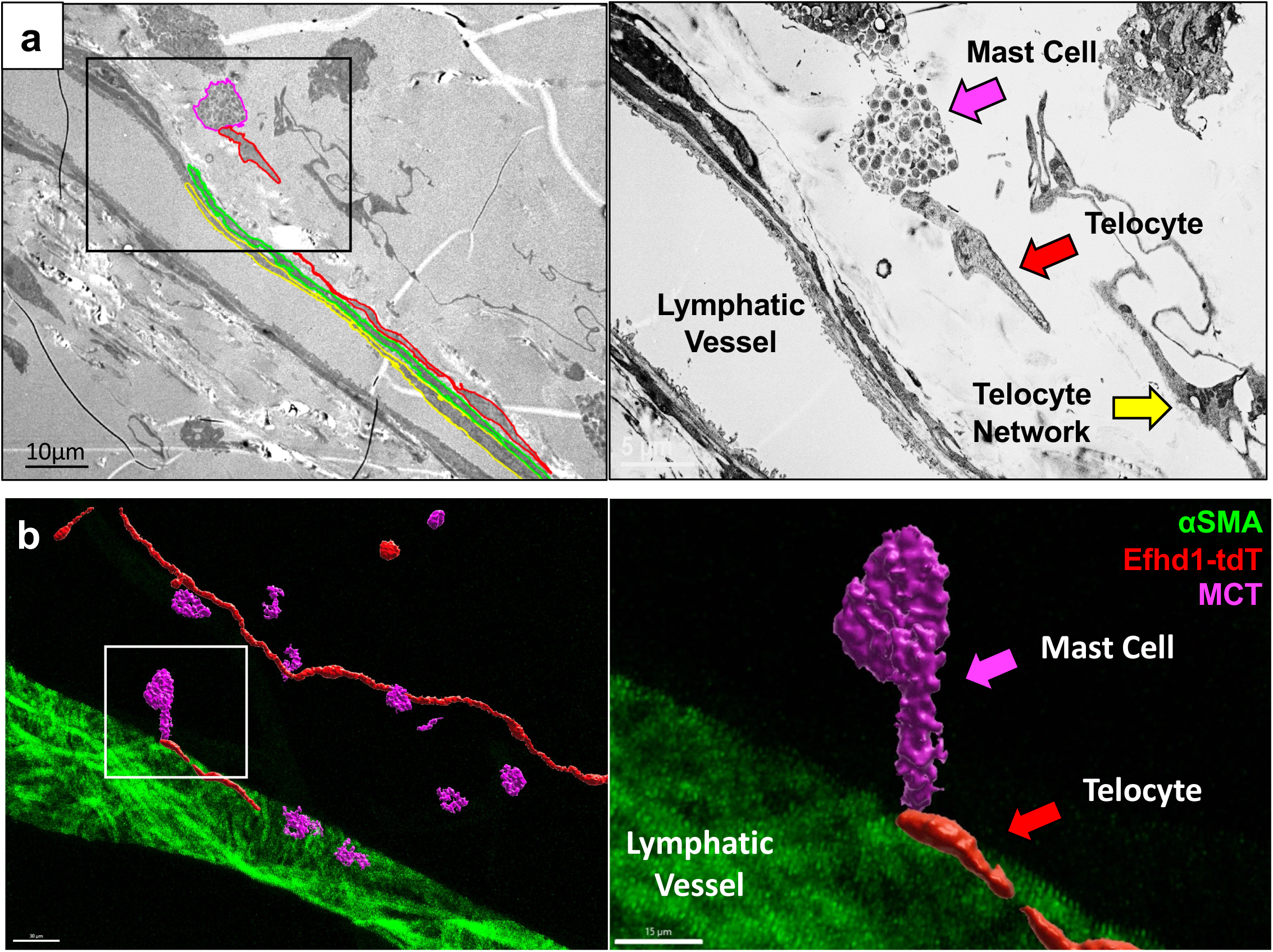
TEM and WMIFM imaging of pre-PLV mast cells physically integrated into telocyte arrays. **a** TEM was performed on the PLVs described in Fig. 2 and a representative x1000 image is shown highlighting the endothelium (yellow) and smooth muscle (green) layers, cells with telocyte ultrastructural morphology (red), and a mast cell (magenta). A high power x5,000 image of the boxed region is shown highlighting the telocyte-like cell (red arrow) physically integrated into a mast cell (purple arrow), and remnants of a network of telocyte-like cells (yellow arrow). **b** PLV with tdT^+^ telocyte-like cells (red) were labelled with fluorescent antibodies against αSMA (green) and mast cell tryptase (MCT; magenta) for WMIF confocal microscopy, and a representative 3D rendering is shown. Note several mast cells are tethered to the tdT^+^ telocyte-like cell network (**Supplemental Video 1**). The magnified image of the boxed ROI illustrates the physical integration, as the distance between the mast cell and tdT^+^ telocyte-like cell is less than 500nm.

To provide additional confirmation that the tdT^+^ cells are telocytes, we reanalyzed our published scRNAseq data on murine PLV-associated cells (*32*) to identify the *Pdgfr* ^+^/*Cd34*^+^ population known to contain telocytes and adventitial fibroblasts (*9*), which produced a uniform manifold approximation and projection (UMAP) of these closely related populations (Fig. 4a). To characterize their transcriptome distinctions and similarities, we generated UMAPs of known marker genes selectively expressed in either one cell type or both telocytes and fibroblasts. The results confirmed that *Myoc, Efhd1, Pla1a, Dpp4, Pcsk6,* and *Ppp2r2b* genes (*9*) are preferentially expressed in PLV-associated telocytes (Fig. 4b). The results also confirmed that: i) the pan marker genes *CD34, Pdgfra,* and *Pdpn* have equivocal expression in both PLV-associated telocytes and fibroblasts, ii) both cell types produce EMC gene products including *Col1a1, Fn1,* and *Lama4*, and iii) *Comp, Dkk3* and *Angpt2* are specific marker genes for PLV-associated fibroblasts. We also performed a cellular trajectory analysis of the scRNAseq data from WT and TNF-tg PLVs using Monocle 3, which predicts that telocytes have the potential to differentiate into fibroblasts (Fig. 4c top). Moreover, we found that there was a 47.7% decrease in the proportion of telocytes relative to fibroblasts in TNF-tg mice relative to their WT littermates (Fig. 4c bottom).

**Fig. 4.**
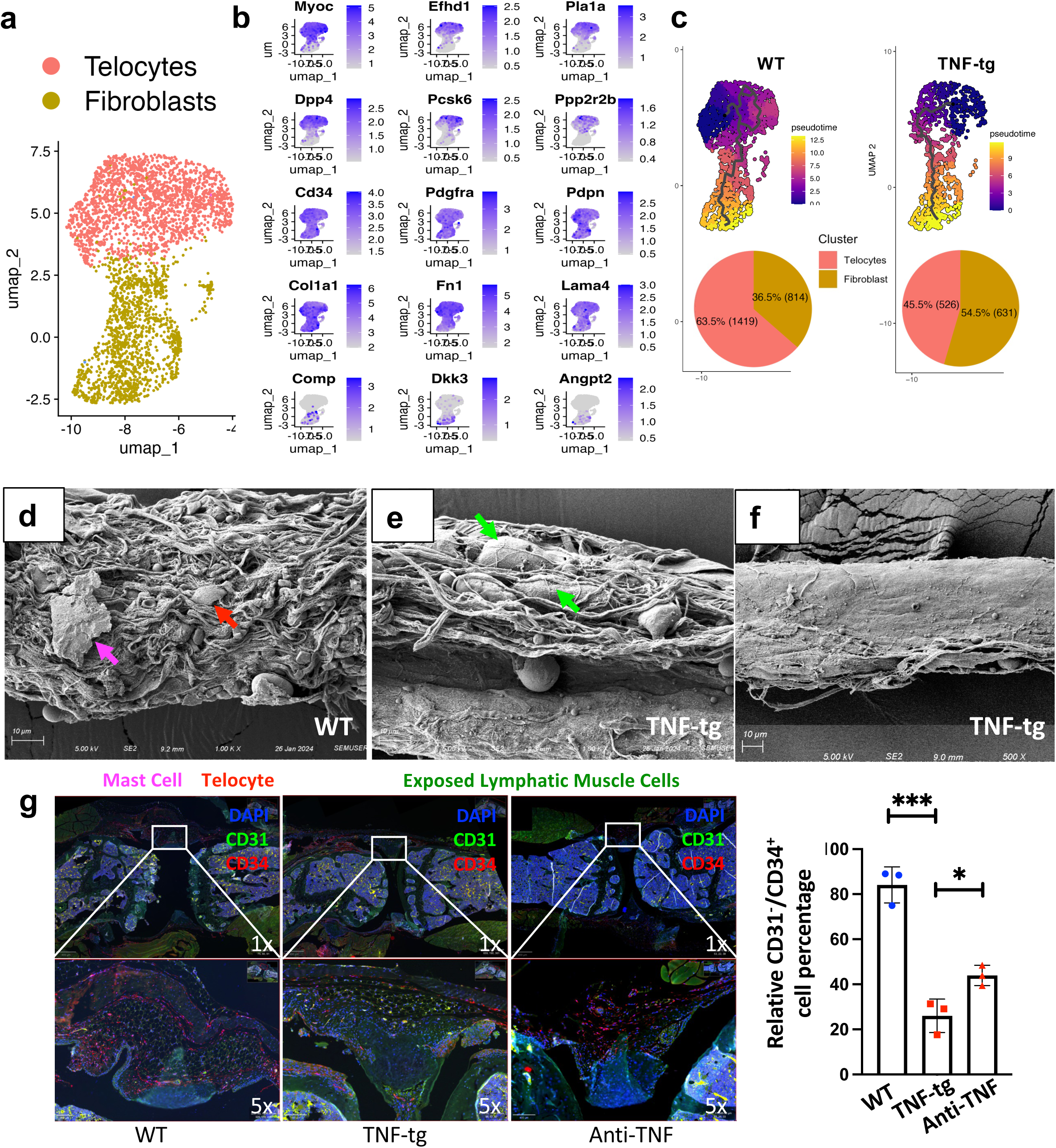
Transcriptome confirmation of PLV-associated telocytes and the loss of PLV extracellular matrix (ECM) and telocytes in TNF-tg mice. **a** Published scRNAseq data on PLV-associated cells from mice^47^ were reanalyzed to identify the Pdgfrα^+^/Cd34^+^ population known to contain telocytes and adventitial fibroblasts^13^, and a color-coated UMAP of these populations associated with WT PLV is presented to demonstrate their proportions. **b** UMAPs of known marker genes selectively expressed in one cell type or both telocytes and fibroblasts are shown to further define these populations. **c** Monocle-3 analysis of the scRNAseq data was performed to generate UMAPs with pseudotime predictions (heat maps), hypothesized telocyte differentiation into fibroblasts (lines within the UMAPs), and the 52.3% decrease in the telocytes:fibroblasts ratio in TNF-tg mice relative to their WT littermates (p>10-15 via Fisher’s test). **d** Representative x5000 SEM image of a PLV from a WT mouse with intact ECM structure, associated mast cell (magenta arrow), and an apparent telocyte cell body (red arrow) embedded in the ECM. Representative x5000 SEM images of PLVs from TNF-tg mice, **e** one with partial ECM loss and exposed LMCs (green arrows), and **f** one with extensive ECM and LMC loss. **g** Immunohistochemistry was performed on knees sections from WT, TNF-tg mice receiving Placebo or Anti-TNF treatment from a prior study^26^, and representative fluorescent microscopy images are shown with quantification of the CD31^-^/CD34^+^ cells as a percentage of synoviocytes (n=3; p<0.001 t-test).

To elucidate synovial peri-lymphatic telocyte loss in TNF-tg mice, we performed scanning electron microscopy (SEM) of PLVs and immunohistochemistry (IHC) on knee synovium. SEM of WT PLVs revealed apparent telocyte cell bodies embedded within a very rich ECM structure, and associated mast cells on top of this ECM (Fig. 4d). In contrast, SEM of TNF-tg PLVs displayed varying degrees of degeneration from partial ECM loss and exposed LMCs (Fig. 4e) to extensive ECM and LMC loss (Fig. 4f), which is consistent with their contractility defects observed *ex vivo* (*25*). Consistent with the loss of CD31^−^/CD34^+^ telocytes observed in RA synovium determined by IHC (*37*), we also observed a significant decrease in CD31^−^/CD34^+^ telocyte-like cells in the synovium of TNF-tg mice receiving placebo treatment compared to WT, and their partially recover from the TNF-tg mice receiving 6 weeks of anti-TNF treatment (*26*) (Fig. 4g).

To confirm that the CD31^−^/CD34^+^ synoviocytes are tdT^+^ telocytes in our transgenic mouse models, we performed flow cytometry, scRNAseq, and immunohistochemistry on lower limb joint tissues from tamoxifen-treated *Efhd1*-CreER^+/-^ x Ai9^+/-^ and *Myoc*-CreER^+/-^ x Ai9^+/-^mice (Fig. 5a). Flow cytometry of knee and ankle synoviocytes revealed consistent profiles of tdT^−^ and tdT^+^ cells (Supplemental Fig. 2). The scRNAseq analysis of the flow sorted tdT^+^ synoviocytes identified 13 distinct cell clusters classified by marker gene expression (Supplemental Fig. 3a-b). To better characterize telocytes in the synovial cells population we re-clustered the mesenchymal and stromal cell compartments (Supplemental Fig. 5c-e). The clusters in the synovium closely resembled the telocyte and fibroblast clusters in lymphatics (Fig. 5b). Specifically, only cells in the upper portion of the UMAP express *Efhd1* and *Pla1a* together with the conventional non-specific telocyte markers *Cd34, Pdgfra, and Pdpn* (*37, 38*), which are broadly expressed throughout the UMAP (Fig. 5b). In contrast to its selective expression in PLV-associated telocytes, *Myoc* is broadly expressed in tdT^+^ synoviocytes with other ECM genes including: *Col1a1, Fn1,* and *Lama4* (Fig. 5b). Interestingly, cells in the lower portion of this UMAP express markers associated with RA fibroblast-like synoviocytes (FLS) (*39, 40*) such as, *Cdh11*, *Comp,* and *Dkk3* (Fig. 5b). Consistent with our observation that the *Efhd1^+^/Pla1a^+^*tdT+ synoviocytes are telocytes, these cells are also negative for the endothelial marker gene *CD31/PECAM1* (*41*), and the mesenchymal stromal cell marker *Cd90/Thy-1* (*42*) (Fig. 5b and Supplemental Fig. 4). However, they are also negative for *FoxL1* (*43*) and *Ano1* (*44*) (Fig. 5b), previously reported telocyte genes identified in gastrointestinal interstitial cells of Cajal (ICC) (*35*).

**Fig. 5.**
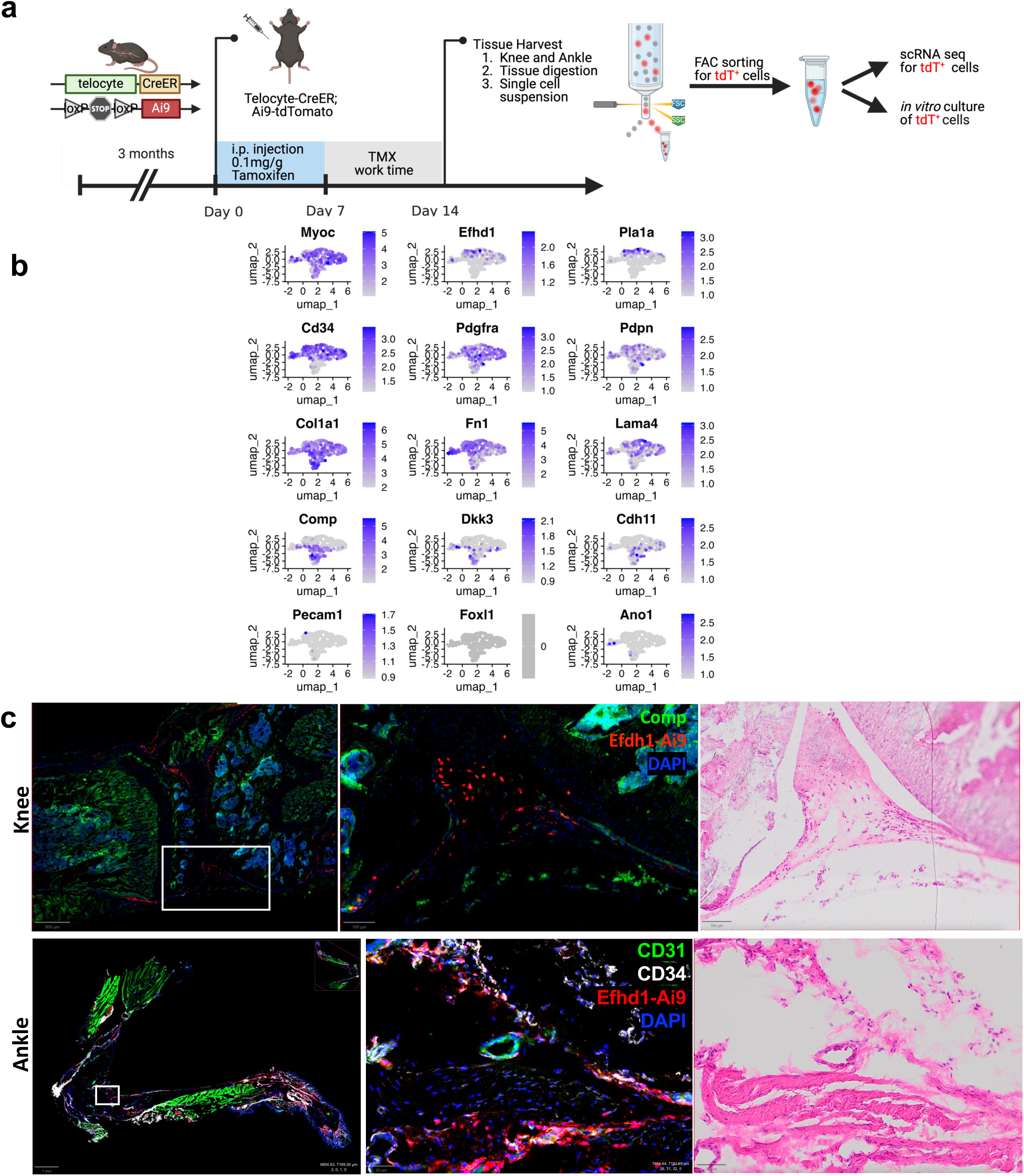
Similarities between synovial and PLV-associated tdT^+^ cells with a telocyte phenotype and their distinctions from fibroblasts. **a** Experimental workflow showing tamoxifen treatment of Efhd1-CreER^+/-^x Ai9^+/-^ mice followed by tissue harvest, cell isolation, and subsequent analyses. **b** Flow cytometric analysis of knee and ankle synovial tissues revealing distinct tdT^high^ and tdT^low^ populations (red arrows). **c** UMAPs showing expression of telocyte and fibroblast marker genes in knee synovial cells, matching the expression patterns observed in Figure 3b. **d** Representative immunofluorescence images of knee and ankle sections from tamoxifen-treated Efhd1-CreER^+/-^ x Ai9^+/-^ mice stained for Comp (green) and DAPI (blue), demonstrating distinct spatial localization of telocytes near the meniscus attachment sites while fibroblasts (Comp^+^) occupy peripheral regions. Images at 1x magnification with 5x magnification of boxed region are shown with parallel H&E bright field images.

Based on these scRNAseq results, we hypothesized the existence of tissue-specific telocyte subsets and performed IHC studies on various tissues from tamoxifen-treated *Efhd1*-CreER^+/-^x Ai9^+/-^ and *Myoc*-CreER^+/-^ x Ai9^+/-^ mice. The results from knee and ankle cryosections confirmed that tdT^+^ synoviocytes are CD31^−^/CD34^+^ cells with telocyte morphology and are distinct from Comp^+^ FLS and CD31^+^/CD34^+^ endothelial cells (Fig. 5c). Given that telocytes are also known to express c-Kit (*35*), a critical gene for mast cells, we examined if c-Kit is also required for telocyte development and survival. Thus, we also included joints from mast cell deficient *Kit^W-sh/W-sh^* (cKit^−/-^) mice and confirmed that they have similar levels of synovial CD31^−^/CD34^+^ telocyte-like cells compared to WT mice (Supplemental Fig. 5), which affirms that lymphatic disfunction in these animals (*31*) is likely due to the loss of mast cells and not loss of telocytes that also express c-Kit. Additionally, we found tdT^+^ telocyte-like cells in tendon, skin hair follicles, and within the ganglion cell layer of the retina, but no tdT^+^ cells were found among the c-Kit^+^ ICC in intestine (Supplemental Fig. 6). Therefore, despite the absence of definitive markers or functional assays to prove telocyte identity, the preponderance of our ultrastructural, histologic, and gene expression data show that the *Efhd1*-CreER^T2^ and *Myoc*-CreER^T2^ mice target tamoxifen-induced gene expression to telocyte subtypes in some but not all tissue types.

### *In vivo* Depletion of Telocytes Decreases Lymphatic Function and Exacerbates Inflammatory-Erosive Arthritis

To test the hypothesis that in vivo depletion of *Myoc* and *Efhd1* expressing telocytes in the synovial lymphatic system are critical for lower limb lymphatic function in mice, we generated an inducible depletion model by crossing the telocyte-specific CreER mice with DTA^floxed^ mice, followed by tamoxifen treatment to induce diphtheria toxin-α *in vivo* depletion of the gene targeted cells as outlined in Figure 6a. To estimate *in vivo* depletion efficiency in these models, we performed WMIFM on PLVs and IHC knee joints from tamoxifen-treated double and single (control) transgenic mice and assessed CD31^−^/CD34^+^ cells. WMIFM of PLVs demonstrated a dramatic DTA elimination of telocytes on the vessel surface and the arrayed telocytes in the adjacent adipose tissue (Fig. 6b). IHC analysis of the knees confirmed a ∼60% reduction in the number of CD31^−^/CD34^+^ synoviocytes (Figs. 6c-d). We repeated this experiment to assess telocyte depletion effects on lymphatic drainage. Cohorts of single and double transgenic mice underwent NIR-ICG imaging prior to tamoxifen treatment and 6hrs following the last tamoxifen injection. We observed no tamoxifen effects on DTA^f/-^ mice contrasted with a significant reduction of ICG clearance in the tamoxifen-treated double transgenic mice (Figs. 6e-f). These findings support our hypothesis that synovial lymphatic system telocytes regulate synovial lymphatic clearance, although these results are confounded by our inability to assess systemic effects and DTA deletion of non-telocytes (e.g. fibroblasts that differentiated from telocytes after tamoxifen treatment).

**Fig. 6.**
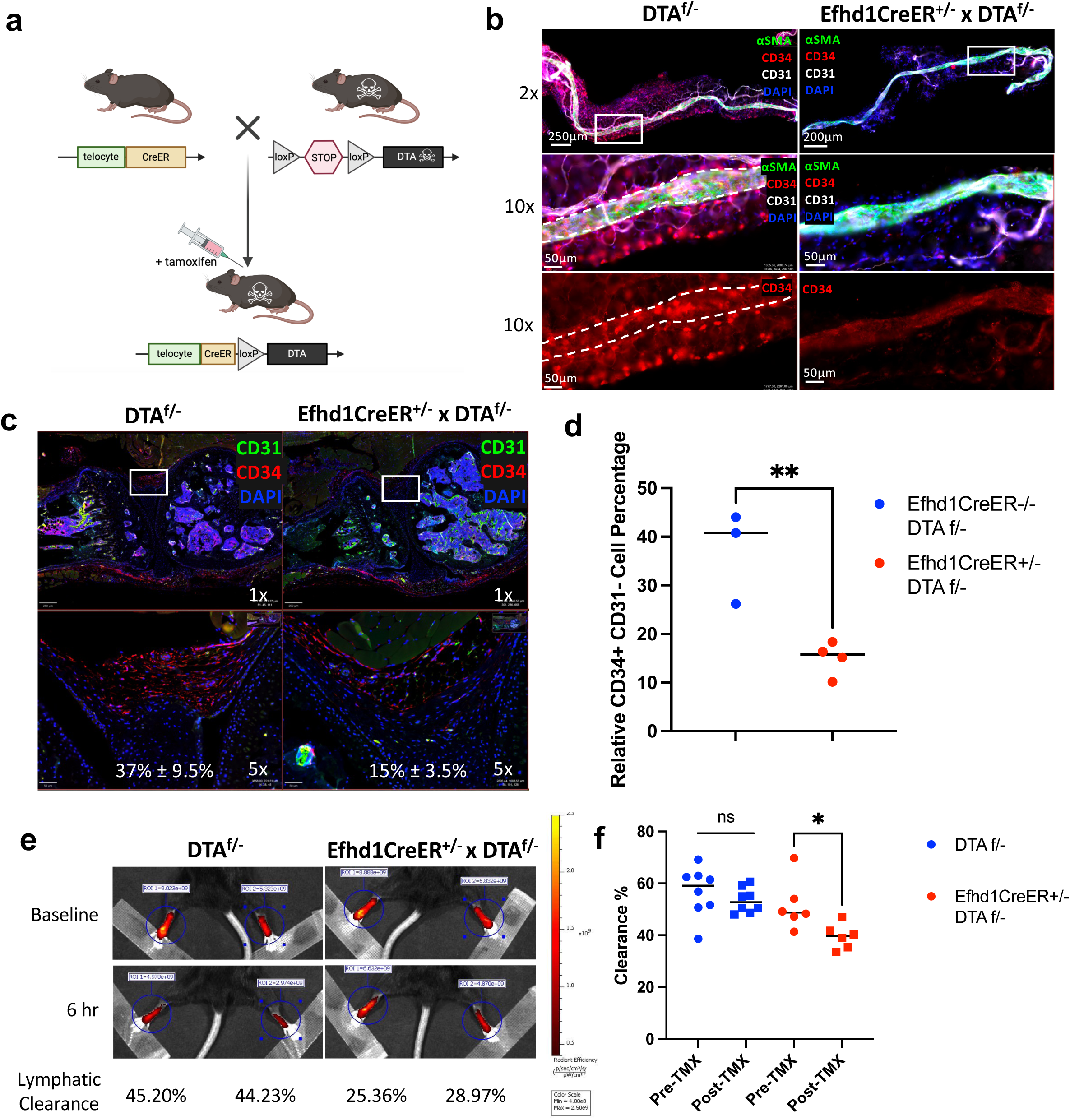
In vivo depletion of telocytes decreases lymphatic function. **a** Schematic illustration of the tamoxifen-induced telocyte deletion model generated by crossing Efhd1-CreERT2 mice DTA-floxed mice. The efficiency of in vivo telocyte deletion following tamoxifen treatment was assessed via **b** WMIFM of PLVs and **c** IHC of knee synovium with quantification of the relative telocyte percentage as described in Fig. 3, from the double transgenic mice and their DTA^f/-^ littermates. **d** Histomorphometric quantification of relative telocyte percentage (n=3; p < 0.5* or 0.001*** vs. Placebo treated TNF-tg via t-test). **e** Lymphatic clearance in tamoxifen-treated Efhd1-CreERT2^+/-^ x DTA^f/-^ mice and their DTA^f/-^ littermate controls was assessed via NIR-ICG imaging, and representative IVIS images of the ICG injected into the footpads at the time of injection (Baseline) and 6hrs later are shown with the heatmap of the signal intensity and % lymphatic clearance for each foot. **f** Data on the % lymphatic clearance of the cohort before and after tamoxifen injection are presented for each mouse with the mean for the group (n≥6; p<0.05 via t-test). Similar results were obtained in experiments with Myoc-CreERT2^+/-^ x DTA^f/-^ mice (data not shown).

To assess the effects of lymphatic dysfunction following telocyte deletion on acute joint inflammation, we utilized the zymosan-induced arthritis (ZIA) model (*45*) as outlined in Figure 7a. Longitudinal ultrasound confirmed a 10-fold increase in synovial volume following knee injection with zymosan vs. PBS controls (Fig. 7b-d). Moreover, zymosan-induced 3.2-fold greater synovitis in telocyte depleted mice, and while this acute synovitis commenced resolution in WT mice after 7 days (Fig. 7c), the increased synovial volume was unabated at day 14 in telocyte depleted mice (Fig. 7d). Histology of the knee joints corroborated the imaging results demonstrating increased pannus tissue in knees with ZIA from mice with depleted telocytes (Figs. 7e-f). Remarkably, the histology also revealed resorbed cortical bone surfaces in DTA and zymosan groups (Fig. 7e). To assess telocyte depletion and ZIA effects on focal erosions and osteoclast numbers, we performed histomorphometry and micro-CT analyses on these femurs. The results showed increased numbers of tartrate-resistant acid phosphatase (TRAP) positive osteoclasts on resorbed cortical bone surfaces in some of the femurs from zymosan treated mice (Supplemental Fig. 7). Surprisingly, we also observed large numbers of osteoclasts in some femurs from telocyte-depleted mice injected with PBS. Micro-CT further demonstrated that acute ZIA failed to induce focal erosions in the knees of WT mice within 14 days, whereas it induced extensive erosions in the patellar groove of telocyte-depleted mice at this time point (Fig. 7g-h). Together with our lymphatic readouts, these data demonstrate that telocyte depletion reduces lymphatic function, and upon challenge with acute inflammatory arthritis, leads to more persistent inflammation, exacerbated synovitis, and rapid focal cortical bone loss.

**Fig. 7.**
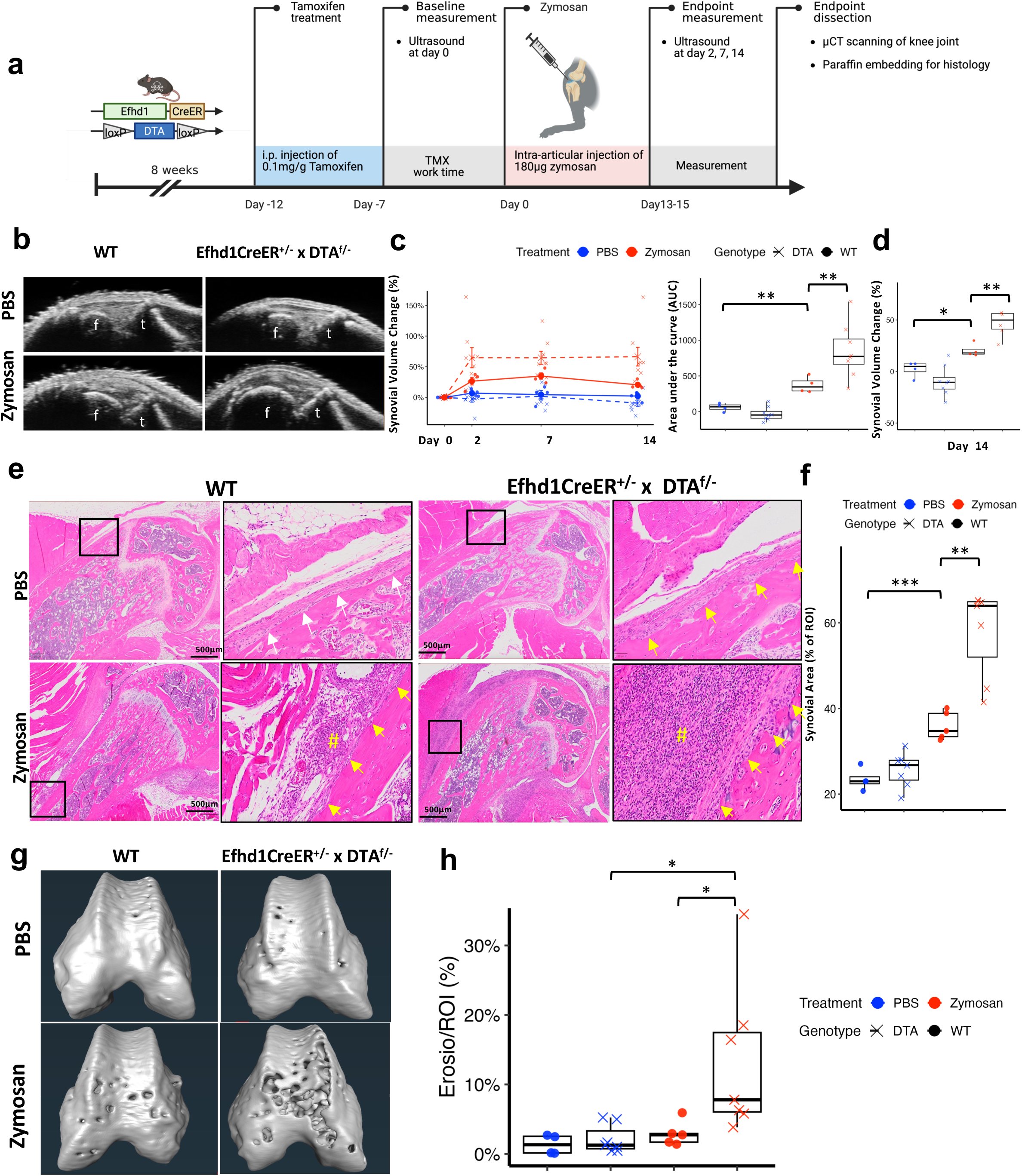
Telocyte depletion increases severity and duration of synovial inflammation from zymosan induced arthritis (ZIA). **a** Experimental design and workflow in which 8-week-old female WT and Efhd1-CreERT2^+/-^ x DTA^f/-^ mice received tamoxifen treatment to introduce telocyte depletion in DTA double-transgenics. Knee injections of zymosan or PBS were given at 10-weeks of age followed by knee ultrasound pre-knee injection and on days- 2, 7, and 14 post-knee injection. Mice were euthanized on day-15 and knee joints were harvested for histology. **b** Representative day-14 knee ultrasound images of the synovial volume between the femur (f) and tibia (t) that were quantified longitudinally (**c** AUC values with mean +/- SD; p<0.01, ANOVA) and prior to sacrifice on day-14 time (**d**; **p ≤ 0.01 via ANOVA). **e** Representative H&E-stained joint histology obtained at 10x with magnified regions of interest (box) highlighting non-resorbed cortical bone surfaces in PBS WT (white arrows) vs. resorbed cortical bone surface in all other groups (yellow arrows), and the ZIA pannus tissue (#). **f** Quantification of the synovial area is reported as the percentage of ROI, as described in Methods. Data are presented with the mean +/- SD (**p<0.01via ANOVA). Similar results were obtained in experiments with Myoc-CreERT2^+/-^ x DTA^f/-^ mice (data not shown). **g** 3D renderings of the distal femur micro-CT scans, with quantification of the focal erosions(**h;** *p ≤ 0.05 via ANOVA). Data are presented for each femur with mean +/- SD.

### Telocyte vs. fibroblast responses to osmotic stress in vitro

Based on literature demonstrating Ca^++^ signaling in telocytes (*46*), Efhd1 regulation of the mitochondria calcium uniporter (MCU) (*47*) and mitoflash (*48*), and our discovery of the peri-PLV telocyte network (*49*), we hypothesized that telocytes within the synovial lymphatic system function as sensors of joint edema that trigger lymphatic muscle cell contractions in resting joint-draining collecting lymphatic vessels (cLV) via intracellular Ca^++^ signaling through mitochondria. To test this in vitro, we developed a primary cell model in which PLVs from tamoxifen-treated *Efhd1*-CreERT2 x Ai9 mice were grown as explants to generate co-cultures of tdT^+^ telocytes and tdT^−^ fibroblasts, which were then loaded with Fluo4 and Ca^++^, Hoechst stained, and challenged with an osmotic shock from 100mM sucrose as outlined in Figure 8a. The results demonstrated that a greater proportion of telocytes undergo an intracellular Ca^++^ flux in response to this osmotic shock than fibroblasts (Fig. 8b). To assess if this intracellular Ca^++^ flux is associate with increased mitoflash, which is a mixed signal consisting of both transient bursts in superoxide production coupled to a modest pH alkalinization and depolarization of the mitochondria inner membrane (MIM) (*50*), we performed osmotic shock experiments with telocyte:fibroblast co-cultures. The results demonstrated high levels of superoxide in the mitochondria of tdT^+^ telocytes, but not in tdT^−^ fibroblasts, following stimulation with antimycin (positive control) and osmotic shock with sucrose, as detected by MitoSOX Green (Figs. 8c-e and Supplemental Video 2-4). This finding demonstrates the remarkable specificity of this signaling pathway. In contrast, recombinant adenovirus transfection of the co-cultures with circularly permuted yellow fluorescent protein (mt-cpYFP) that detects pH changes (*51*) revealed flashes in both tdT^+^ telocytes and tdT^−^ fibroblasts that were modestly increased following stimulation with pyruvate (positive control) and osmotic shock with sucrose (Supplemental Videos 5-7).

**Figure 8.**
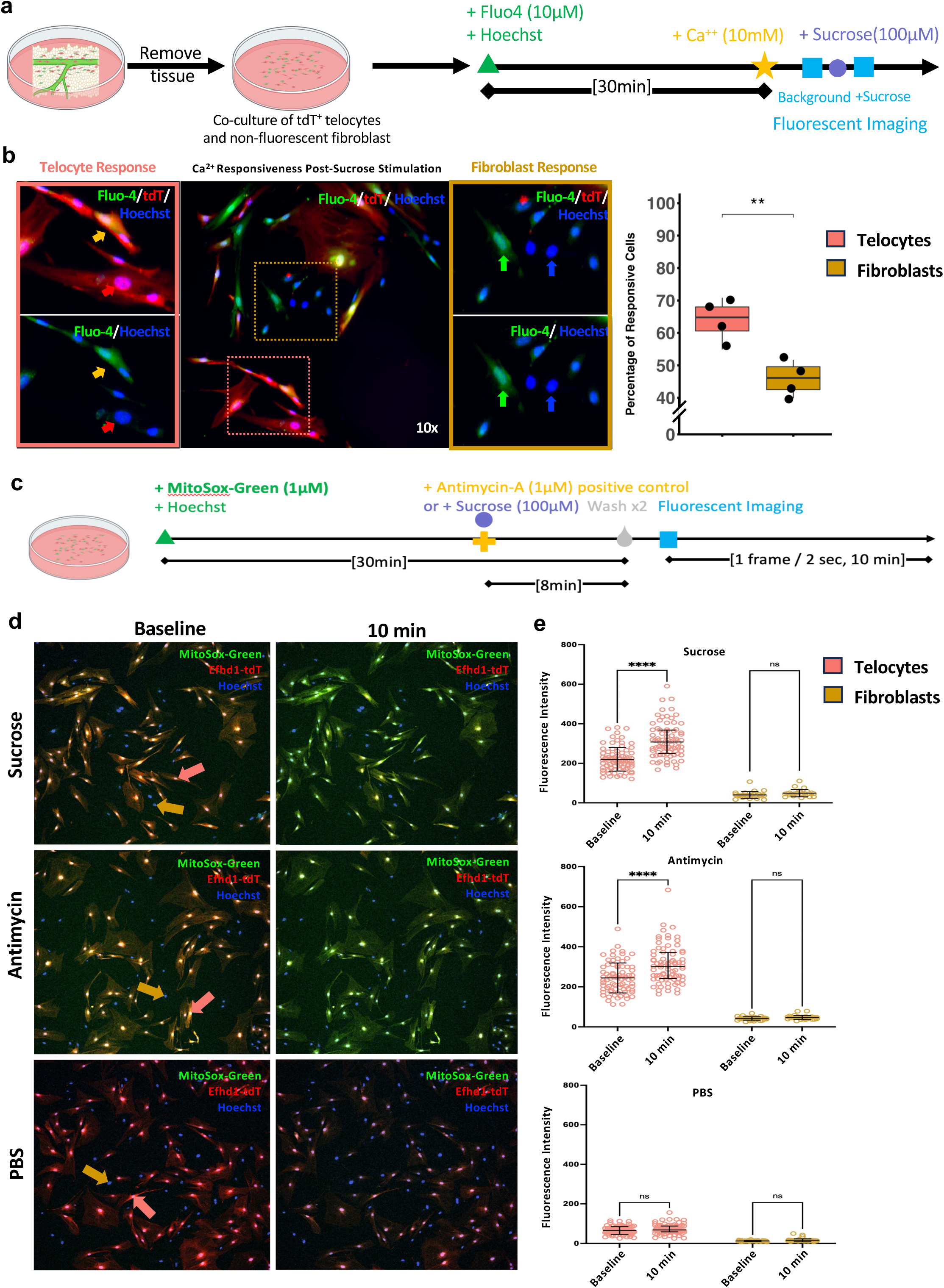
Telocytes are more sensitive to osmotic pressure changes than fibroblasts and uniquely respond by increasing mitochondrial superoxide. **a** Schematic illustration of plated PLVs from tamoxifen treated Efhd1-CreER^+/-^x Ai9^+/-^ mice to establish primary co-cultures of tdT^+^ telocytes and tdT^-^ fibroblasts. For osmotic shock studies the cells were preloaded with Hoechst, Ca^++^, and Fluo-4, and incubated with 100µM sucrose followed by real-time fluorescent microscopy. **b** Representative 10x fluorescent images of a co-culture post-sucrose stimulation are shown to illustrate Ca^++^ responsiveness seen as green fluorescence (center). The ROIs (dotted boxes) are magnified in the flanking images to illustrate four cell types: non-responsive telocytes (red arrows), responsive telocytes (orang arrows), non-responsive fibroblasts (blue arrows), and responsive fibroblasts (green arrows) with (top) and without (bottom) the superimposed tdT channel. The response to 100 µM sucrose was quantified as the % Fluo-4 positive telocytes or fibroblasts, and the data are plotted with the mean +/- SD (** p<0.01, t-test, each data point represents the mean value of 4 images from a single chamber from an independent experiment). **c** Schematic illustration of the co-culture incubated with MitoSOX green and Hoechst 30 minutes prior to real-time fluorescent microscopic imaging. Eight minutes prior to imaging the co-cultures were challenged with 100 µM sucrose, 1 μM antimycin (positive control), or PBS (negative control). **d** Representative 10x fluorescent images of the three treatment groups at the beginning of the imaging session (Baseline) and 10 minutes later are shown to illustrate accumulation of MitoSOX green fluoresce over time only in the telocytes (pink arrows) but not in the fibroblasts (orange arrows). **e** Analysis of MitoSOX green fluoresce intensity over time in telocytes versus fibroblasts in three treatment groups. Data are presented as the MFI within the borders of a single nucleus with means +/- SD of the group at each time point (**** p<0.0001, one-way ANOVA).

### Telocyte vs. fibroblast invasiveness through ECM in vitro

Efhd1 inhibition of the MCU and the Hippo signaling pathway maintains cells in a sessile state, and its loss in tumor cells is associated with increased metastatic potential (*47*). Thus, we hypothesized that *Efhd1*-expressing telocytes possess a non-invasive phenotype compared to FLS. To test this, serum-starved tdT^+^ telocyte and tdT^−^ fibroblasts co-cultures were added to Matrigel transwell inserts and allowed to migrate towards culture media ± 5 ng/mL TGF-β1 as the chemoattractant in the lower chamber prior to fixation, Hoechst staining, and fluorescent microscopy as illustrated in Figure 9a. The results demonstrated that the fibroblasts invaded further into the Matrigel under both conditions (Figs. 9b-c).

**Figure 9.**
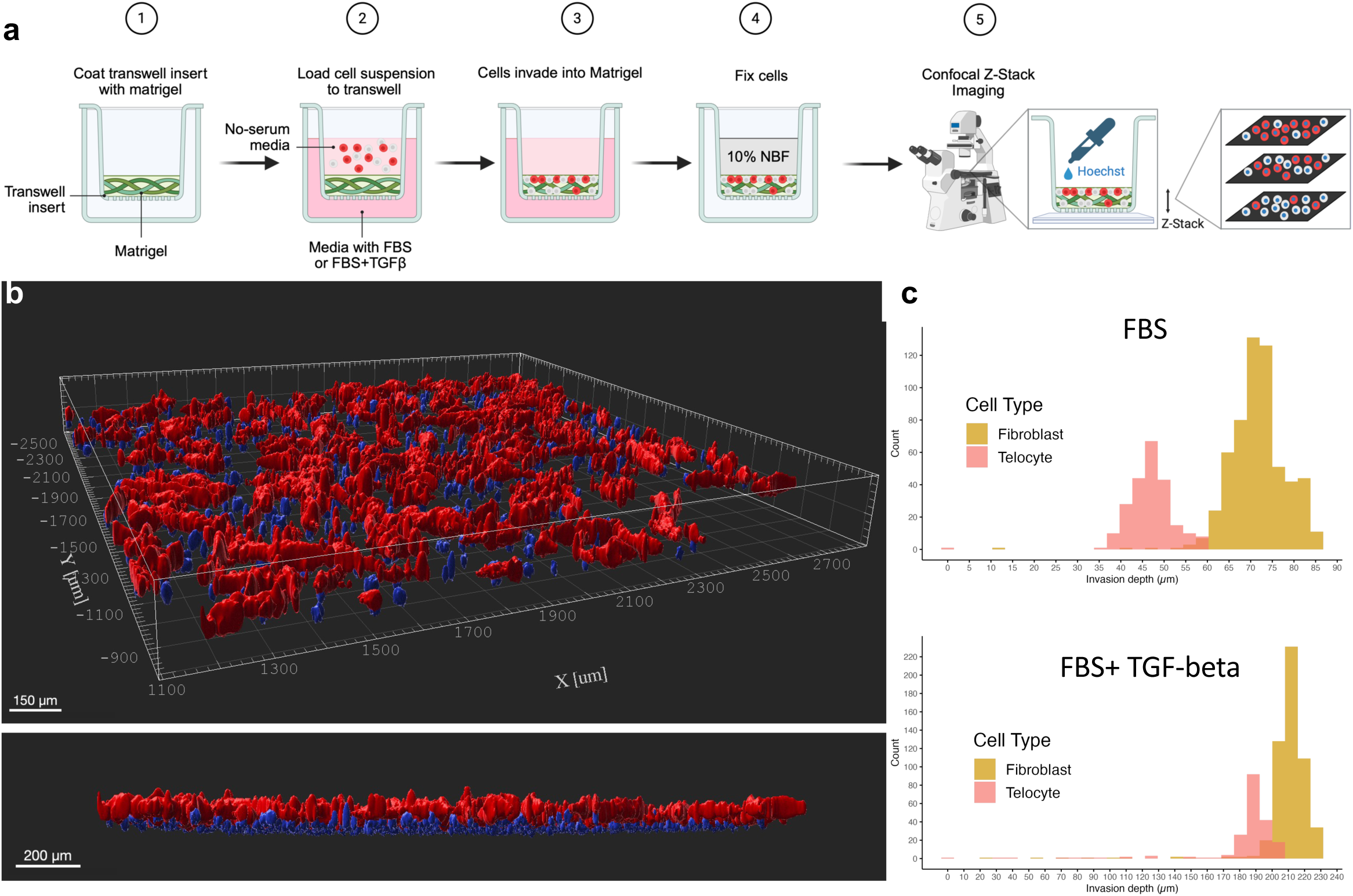
Telocytes are less invasive than fibroblasts. **a** Schematic overview of the Matrigel invasion assay workflow. Telocytes and fibroblasts were co-cultured on transwell inserts (1), serum-starved (2), and placed above Matrigel-containing chambers with serum-free media in the upper compartment and 10% FBS ± 5 ng/mL TGF-β as the chemoattractant in the lower chamber (3). After a 24-hr invasion period (4), the inserts were fixed in 10% neutral buffered formalin and stained with Hoechst prior to imaging (5). **b** Representative 3D reconstructed confocal images of co-cultured telocytes (red) and fibroblasts (blue nuclei) invading into Matrigel. Top and side views illustrate the relative positioning and invasion depth of each cell population. **c** Quantification of invasion depth. The penetration distance of individual Hoechst-labeled nuclei was measured throughout the Matrigel and is presented as histograms for each condition. Data shown are representative of three independent experiments..

### Telocyte to fibroblast transdifferentiation in vitro

Bioinformatic trajectory analysis of the UMAP of peri-PLV cells predicts that telocytes differentiate into fibroblasts (*49*), and this differentiation is predicted to occur with increasing ECM production (Supplemental Fig. 1e). To test this, we obtained FACS purified tdT^+^ synoviocytes from knees of tamoxifen treated *Efhd1*-CreERT2 xAi9 mice (Fig. 5b), cultured them to ensure they retain their tdT^+^ telocyte phenotype over several passages (Supplemental Fig. 8), and then cultured them on collagen-coated CytoSoft® Imaging plates with varying stiffness levels: 0.2 kPa (soft), 8 kPa (intermediate), and 64 kPa (stiff). After 24 hours of culture, the cells were harvested for bulk RNA sequencing and gene expression analyses. The results demonstrated that telocytes grown on a soft matrix maintain their telocyte transcriptome, whereas telocytes grown on a stiff matrix express fibroblast and myofibroblast marker genes (Fig. 10).

**Fig. 10.**
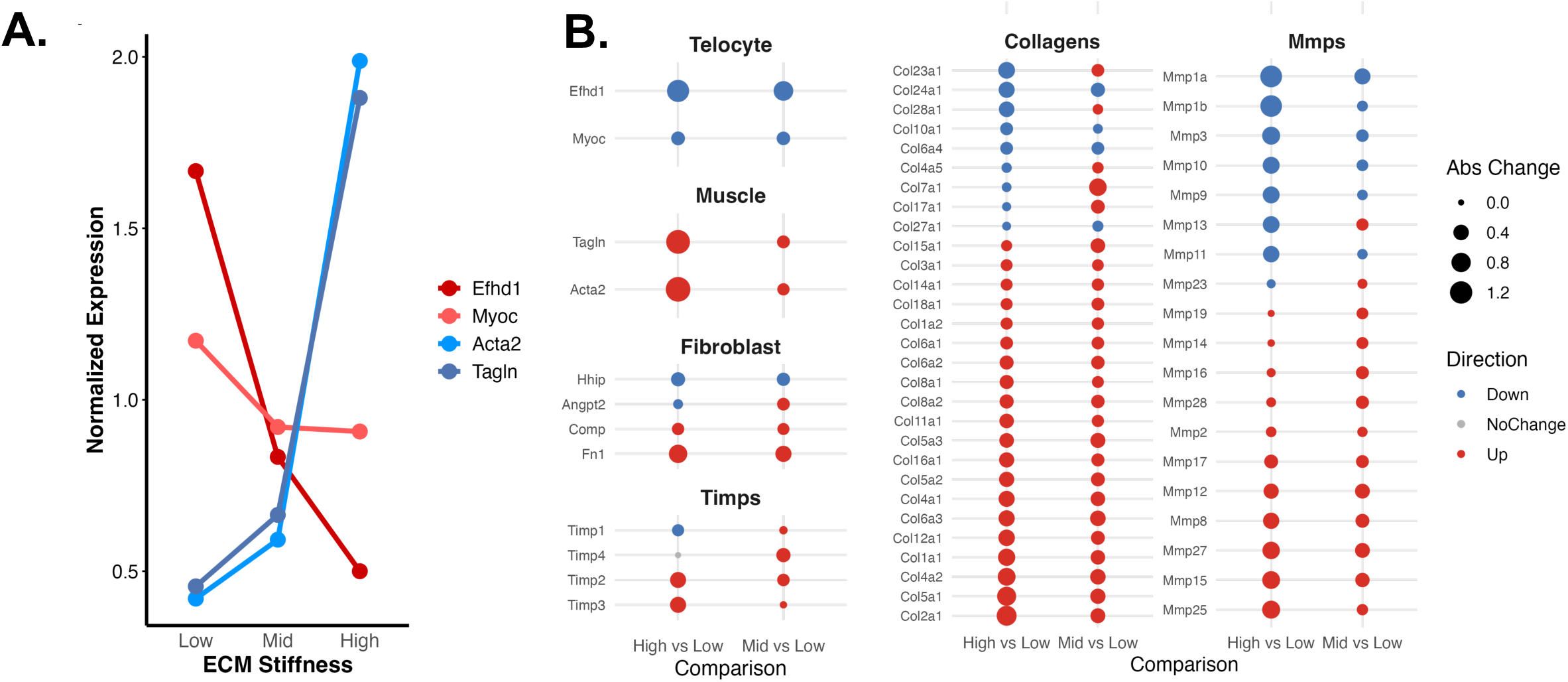
PLV-associated telocytes differentiate into fibroblasts on stiff ECM. Primary cultures of the FACS purified tdT^+^ cells described in Fig. 5 were cultured on collagen-coated CytoSoft® Imaging 24-well plates with varying stiffness levels: 0.2 kPa (soft), 8 kPa (intermediate), and 64 kPa (stiff). After 24hrs of culture the cells were harvested for bulk RNA sequencing and gene expression analyses. **a** Normalized expression of telocyte (*Efhd1* and *Myoc*) and myofibroblast (*Acta2* and *Tagln*) maker genes are graphed as a function of the ECM stiffness. **b** The quantified change in the indicated telocyte, myofibroblast, and fibroblast marker gene expression is presented with changes as indicated by the key. Also presented are the in significantly expressed *TIMP*, *Collagen* and *MMP* genes. All indicated changes are significant (p>0.05).

## DISCUSSION

RA is a complex immune-mediated inflammatory disorder with divergent mechanisms of disease progression (*52, 53*). Recent high-dimensional studies of RA tissues revealed multiple cell types and factors that contribute to chronic inflammation in affected joints (*54, 55*), and this knowledge catalyzed promising treatment strategies for RA patients (*56*). Despite these therapeutic advances, major unmet clinical needs remain for a large fraction of RA patients that fail to achieve remission and ∼30% who are refractory to all current therapies (*52, 53*). This treatment shortfall underscores the need to find novel disease mechanisms and treatment strategies. With this objective in mind, we focused on the autoimmune independent pathologies of inflammatory-erosive arthritis observed in TNF-tg mice (*57*). This research led to the discovery of the synovial lymphatic system (*58*), an interactive network critical for joint homeostasis (*59*), which is dysfunctional in RA and osteoarthritis, and this altered function correlates with disease progression and flare (*13, 14*). Thus, elucidation of the mechanisms that regulate the synovial lymphatic system and its dysfunction during chronic arthritis opens new avenues of investigation for the development of new therapies. Here we demonstrate that DTA-mediated telocyte deletion inhibits normal lymphatic function (Fig. 6) and exacerbates inflammatory-erosive arthritis (Fig. 7), which raises the possibility that these cells might serve as a novel target for treating RA patients who are refractory to all current therapies.

In contrast to the circulatory system, which is responsible for blood flow, and where development, regulation, and repair have been defined in great cellular and molecular detail, many fundamental questions about lymphatics remain unanswered, including the origins of LMCs and cLV maintenance and repair (*3, 60*). Critical questions pertaining to the mechanisms that control cLV contractions also remain unanswered. To address lymphatic contraction mechanisms, Zawieja et al utilized conditional-inducible CreER^T2^ transgenic mice to assess the innate pacemaking potential of mast cells, telocytes, pericytes, fibroblasts and LMCs in cLVs ex vivo (*9*). These studies failed to identify Ca^++^ events in PDGFRα^+^ or c-Kit^+^ adventitial cells that were in phase with cLV contractile activity. These investigators were unable to consistently elicit contractions following optogenetic stimulation via a light-activated cation channel rhodopsin2 (ChR2), indicating that adventitial cells (e.g. telocytes) are not spontaneous pacemakers. However, they did find that MYH11^+^ LMCs exhibited asynchronous diastolic Ca^++^ events that were dynamically modulated by pressure, and these cells also propagated contraction in response to ChR2 photo-stimulated depolarization (*9*). Taken together with other studies (*5, 61*), these results support a long-standing model of LMCs as the intrinsic pacemaker of cLVs (*62*). While these studies largely explain the mechanisms responsible for intraluminal pressure-dependent spontaneous cLV contractions, they do not provide information on the initiating signal for distal tissue LMC contractions in response to edema (e.g. joint effusions).

In retrospect, our finding that ablation of peri-PLV telocytes is associated with lymphatic dysfunction is not surprising, as pacemaking in smooth muscle-invested organs is known to be controlled by this cell type (*6*). Indeed, pioneering studies of gastrointestinal smooth muscle found that pacemaking is initiated by ICC (*63, 64*), and subsequent studies of the lower urinary tract (*65*), gallbladder (*66*), and oviduct (*67*) found similar cells, now generically referred to as telocytes (*35, 36*). Additionally, cells characteristic of telocytes have been identified in LVs (*68*), and prior studies identified methylene blue staining peri-LV CD34^+^/c-Kit^+^ cells with telocyte morphology and telopode-like structures extending to LMCs (*9, 68*).

It also reasons that telocytes are involved in cLV contractions, as LMCs are akin to smooth muscle cells that lack the ionic mechanisms required for action potential regeneration and electrical signals that efficiently propagate between muscle cells (*69*). Thus, LMCs likely rely on a telocyte network for signal conduction. Consistently, telocytes mediate rhythmic electrical activity in other tissues (*70*), which is believed to depend on ion channels for generating pacemaking currents (*71, 72*). Interestingly, electrically coupled cells are typically connected by gap junctions (*68*), and evidence demonstrating this physical link between telocytes and LMCs does not exist (*6*). Studies have also failed to demonstrate functional electrical communication between telocytes and LMCs or a telocyte Ca^++^ clock to drive the rhythmic cLV contractions observed *ex vivo* (*9*). However, it is conceivable, based on our findings, that an intermediary cell provides the depolarizing signals to LMCs to initiate pacemaking, which we posit to be peri-cLV mast cells integrated into telocytes (Fig. 3). Mast cells produce, store, and release various inflammatory and vasoactive mediators (*27, 73*). These mediators influence lymphatic pumping by affecting LMC contractility (*74*), and may be responsible for directly providing the initiating pacemaking signal to LMCs in joint-draining cLV secondary to a Ca^++^ flux through gap junctions from attached telocytes. The Ca^++^ flux generated from telocyte responses to osmotic stress (Figure 8b) and Ehfd1 may play a direct role in calcium signaling via inhibition of the mitochondrial calcium uniporter (*47*), followed by shunting of intracellular Ca^++^ through the telocyte network into mast cells.

Our finding that *Myoc* is a peri-PLV telocyte marker gene is also not surprising given that the extensive ECM (Fig. 4d), and primary functions of this protein are cross-linking ECM proteins and regulating MMPs (*75*). It is highly probable that *Myoc* is not a telocyte-specific gene in the joint (Fig. 5b) since the fibroblasts in these musculoskeletal tissues play a major role in ECM genesis and remodeling. Therefore, since *Efhd1*-CreER^T2^ mice have more selective gene targeting, we conclude that they are superior model for telocyte research. However, mutations in the *Myoc* are known to be the greatest genetic risk factor for glaucoma. Thus, the *Myoc*-CreER^T2^ mice may be a useful model to better understand this disease.

Here we also report the first evidence of tissue-specific telocytes, which includes our failure to detect *Efhd1* and *Myoc* expression in ICC of the gut (Supplemental Fig. 5a). This observation also distinguishes our *Efhd1* and *Myoc* telocyte-targeted CreER^T2^ models from the established *FoxL1*-Cre targeted models (*43, 76*), although all of these mice have transgene targeting in subepithelial telocytes, suggesting overlaps in the tissue specificity of distinct telocytes. Additionally, our scRNAseq data demonstrates that synovial and peri-PLV telocytes do not express *Ano1* (Fig. 5c), which is considered to be the canonical Ca^++^ activated chloride channel in ICC required for pacemaker activity (*71, 77*). Thus, Ca^++^ signaling in telocytes of the synovial lymphatic system must involve different pathways, which might include mitochondria, considering that EFHD1 regulates mitoflash activation that mediates intercellular signaling (*78*) and physically binds to the mitochondrial calcium uniporter to inhibit the Hippo/YAP pathway and cellular invasives (*47*). Taken together with our findings that *Efhd1* expression in telocytes is lost during telocyte to fibroblast/myofibroblast differentiation (Fig. 10), EFHD1 may be a target to stabilize telocytes in arthritis and other emerging fields that propose telocyte transplantation (*79*).

In terms of the well-known fibrosis that is associated with RA and lymphedema, our *in vitro* findings demonstrating that relatively non-motile *Efhd1*^+^ telocytes differentiate into invasive fibroblasts in response to TGF-β and stiff ECM (Figs. 9 & 10) may provide etiologic insights. Thus, prospective lineage tracing studies with tamoxifen-treated *Efhd1*-CreER^T2^ x Ai9 and *Myoc*-CreER^T2^ x Ai9 mice are warranted to see if this telocyte to myofibroblast differentiation also occurs in vivo, especially in disease models.

We present the first report of telocytes within the synovial lymphatic system, coupled with several preliminary findings that must be addressed in future studies. Notable among these limitations are potential off-target effects of our transgenic mouse models when performing systemic tamoxifen delivery experiments. As our models may target cells in tissues that were not studied exhaustively and we confirmed targeting to non-telocytes known to express Efhd1 (e.g. renal tubule cells (*47*) Supplemental Fig. 9), conclusions about telocyte-specific effects need to be tempered. We also report some preliminary results that need to be follow up in future studies. The first is our variable findings of increased osteoclasts in femurs of mice with DTA-deleted target cells (Supplemental Fig. 7), which warrants prospective studies to determine if these preliminary observations are significant. We also conclude that our highly variable NIR-ICG results to assess lymphatic drainage of PBS vs. zymosan injected knees were confounded by effects of multiple intraarticular injections (data not shown), and that faithful measurement of lymphatic drainage from the knee will likely require technical innovations (e.g. assessing large molecule translocation from the joint to iliac lymph nodes).

In summary, we find that telocytes embedded in the ECM around PLVs are lost in TNF-tg mice with inflammatory arthritis, and telocyte networks adjacent to PLVs are integrated into mast cells. Furthermore, telocyte deletion decreases lymphatic function and exacerbates ZIA. Based on these discoveries, we propose a unifying model to explain the functions of telocytes within the synovial lymphatic system (Fig. 11). This model posits three testable hypotheses that are the focus of our future directions. The first is the existence of connected telocyte networks that parallel lymphatic vessels from the synovial lymphatic capillaries to mast cell elaborated cLV, which is supported by our WMIFM images (Fig. 2h) and can be further tested by high-resolution light-sheet microscopy to confirm the tdT^+^ cells in the array are physically associated. The second is that telocytes sense osmotic pressure in joints (e.g. ankle) and signal through the telocyte network to the peri-PLV mast cells that degranulate and initiate LMC contractions in response (Fig. 11b). Experiments to test this will require the generation of double transgenic mice in which the *Efhd1*-CreER^T2^ are crossed to mice with telocyte or mast cell-specific Ca^++^ reporter probes for intravital microscopy studies following ankle stimulations. We also propose that RA-FLS are derived from telocytes that differentiate in response to chronic inflammation and adherence to rigid ECM in inflamed joints (Fig. 11c). This could be tested through lineage tracing studies using tamoxifen-treated *Efhd1*-CreER^T2^ x Ai9 mice, as well as genetic studies on Efhd1 gain and loss of function, which could provide the field with a novel molecular target for treating pauci-immune RA and lymphedema that are refractory to all known treatments.

**Fig. 11.**
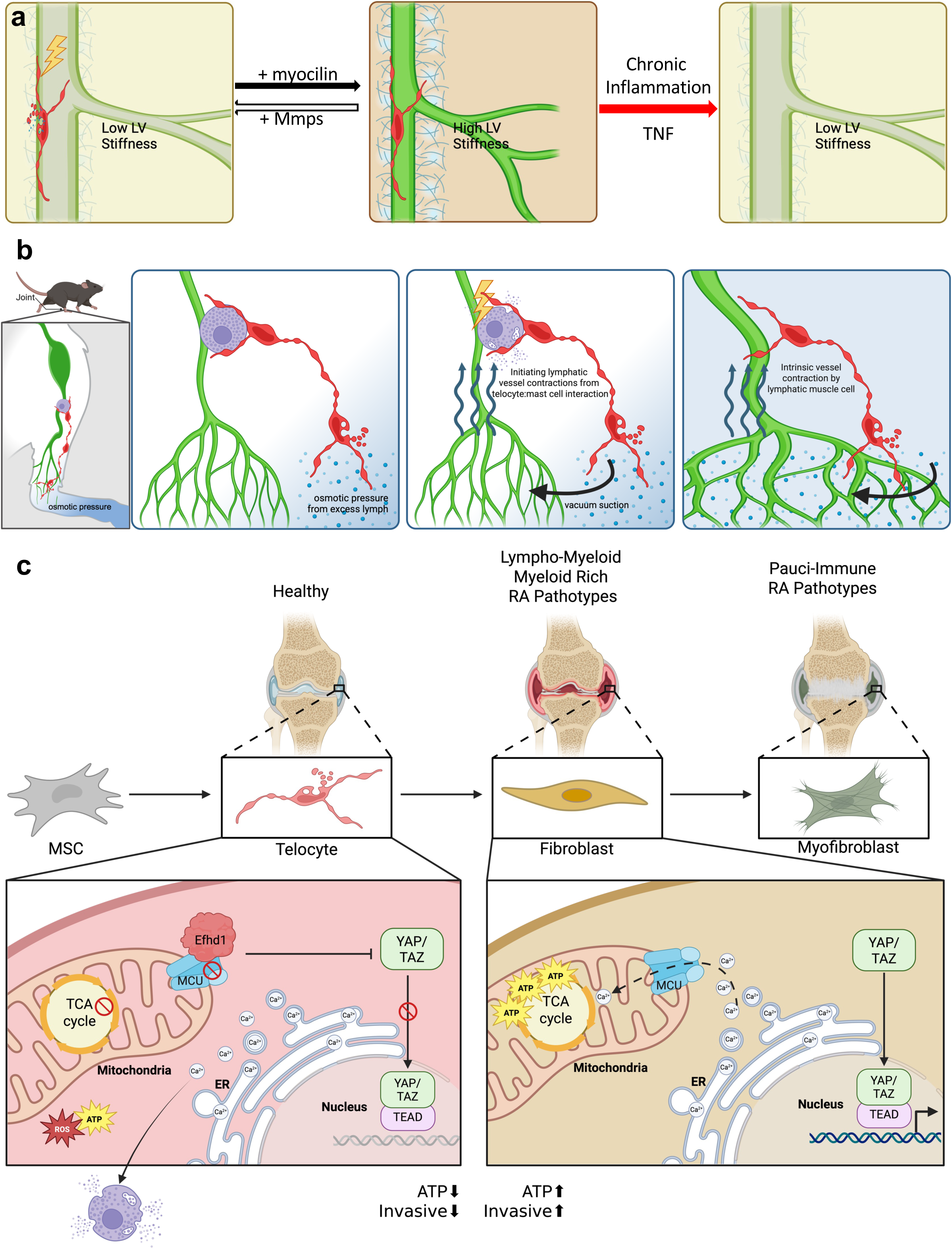
Proposed telocyte networks within the synovial lymphatic system of the lower limb and their regulation of joint homeostasis and lymphatic function. **a** Schematic model of telocyte regulation of PLV stiffness. Telocytes with telopods integrated along PLV monitor stiffness and remodel the ECM. When the PLV is too flaccid, the telocytes produce myocilin to crosslink the collagen and fibronectin in the ECM (black arrow). When the PLV becomes too rigid from ECM accumulation, telocytes release matrix metalloproteases (MMPs) and downregulate tissue inhibitors of MMPs (TIMPs) to degrade the ECM (white arrow). In the absence of telocytes in TNF-tg mice, chronic inflammation perpetuates MMP breakdown of the ECM and dysfunction of flaccid PLV (red arrow). **b** Schematic model of telocyte network regulation of PLV contraction in the mouse lower limb. Telocytes in the synovium monitor extracellular osmotic pressure in joint fluid and are activated upon sensing an excessive threshold (left). Upon activation, an intercellular signal is transmitted through the network of telopod-to-telopod connected telocytes and is ultimately delivered to mast cells adjacent to the joint-draining cLV. This signal triggers mast cell degranulation and release of vasoconstricting factors (e.g. histamine) that initiate LMC contractions in the resting PLV. Subsequent diastole, assisted by ECM tensile force, generates vacuum suction to fill the PLV with lymph from the joint (middle), and the intraluminal shear force triggers intrinsic PLV contractions by LMCs, which then clears the excess interstitial fluid in the joint, returning it to normal osmotic pressure (right). (**c**, left) Schematic model of synovial telocyte differentiation from a mesenchymal stem cell (MSC) and its intracellular biochemistry regulated by Efhd1, which inhibits the mitochondria calcium uniporter (MCU) and cytosolic Ca^++^ import for ATP synthesis from oxidative phosphorylation (*47*). This also inhibits activation of YAP/TAZ in the Hippo pathway, which is critical for cell motility (*47*). We also hypothesize that this shunted Ca^++^ is part of the signal transduction through the telocyte network that culminates with peri-cLV mast cell activation. (**c**, right). Based on our scRNAseq trajectory analysis and in vitro demonstration of Efhd1+ telocyte transdifferentiation into a myofibroblast phenotype on stiff ECM (Fig. 9), we hypothesize that telocytes can lose their Efhd1 expression and adopt a RA-FLS phenotype that includes mitochondria uptake of cytosolic Ca^++^ through the MCU to increase metabolic activity through oxidative phosphorylation. The loss of Efhd1 also leads to a more invasive cell phenotype through increased Hippo pathway signaling. A final differentiation into myofibroblasts is also proposed to occur in end-stage pauci-immune fibrotic RA pannus tissue that is stiffer than normal synovium from extensive ECM synthesis.

## Methods

### Mouse models

All animal research was conducted with approval by the University of Rochester Institutional Animal Care and Use Committee. Three transgenic mouse strains were generated with P2A-CreER^T2^ cassettes, which were knocked into C57BL/6NTac mice via CRISPR/Cas-mediated genome engineering (Taconic Biosciences, Germantown, NY). Cas9 and gRNA were co-injected into fertilized eggs with the targeting vector, and the resulting pups were genotyped by PCR followed by Sanger sequencing analysis to confirm the precise insertion loci (for primers see Table 1). For the Efhd1-CreER^T2^ model, the TAG stop codon in the mouse *Efhd1* gene (NCBI Reference Sequence: NM_028889.3) was replaced by a “P2A-CreER^T2^” cassette (Transcript: 201-ENSMUST00000027472). A synonymous mutation p. R232= (CGG to AGA) was also introduced to prevent the binding and re-cutting of the sequence by gRNA after homology-directed repair. For the Myoc-CreER^T2^ model, the TGA stop codon in the mouse *Myoc* gene (NCBI Reference Sequence: NM_010865.3) was replaced by a “P2A-CreER^T2^” cassette (Transcript: 201-ENSMUST00000028020). For the Pla1a-CreER^T2^ model, the TGA stop codon in the mouse *Pla1a* gene (NCBI Reference Sequence: NM_134102.4) was replaced by a “P2A-CreER^T2^” cassette (Transcript: 201-ENSMUST00000002926).

B6.Cg-*Gt(ROSA)26Sor^tm9(CAG-tdTomato)Hze^*/J (Ai9)(*33*), B6.129P2-*Gt(ROSA)26Sor^tm1(DTA)Lky^*/J (DTA)(*34*) and B6.Cg-*Kit^W-sh^* /HNihrJaeBsmJ (*Kit^W-sh/W-sh^* or cKit^−/-^) mice were purchased from The Jackson Laboratory (Bar Harbor, ME, Ai9 Strain #:007909, DTA Strain #:009669, Kit Strain #: 030764) and maintained at the University of Rochester vivarium. For crossing between the strains, CreER^T2^ conditional inducible mice were used as male breeders, and Ai9 or DTA mice as female breeders. For all *in vivo* longitudinal outcome measures, mice were anesthetized with 1–3% isoflurane. All mice were euthanized with a lethal dose of ketamine/xylazine cocktail (intraperitoneal) followed by cervical dislocation.

### Whole mount immunofluorescent microscopy (WMIFM)

PLVs were harvested and fixed as previous described(*80*). Briefly, fixed PLVs were blocked with 5% normal goat serum (NGS; ThermoFisher Scientific Cat# 50062Z)/1 × TBS/0.3% Triton X-100 for 1 hr at RT, and then incubated with the primary antibodies diluted in 5% NGS/1 × TBS / 0.3% Triton X-100 overnight at 4 °C. For αSMA labeling, primary antibodies include mouse anti-αSMA antibody (AlexaFluor 488 conjugate; ThermoFisher Scientific Cat# 53-9760-82, diluted at 1:100). To validate the mast cell-telocyte interaction, the following primary antibodies targeting mast cells were used in this study: Rabbit anti-human Mast Cell Tryptase IgG (mouse reactivity; Bioss Antibodies Cat# BSM-52533R; 1:100 dilution). To prevent non-specific labeling of Fc receptors (FcR), exclusively secondary antibodies below featuring F(ab) fragments were employed: Anti-rabbit IgG (H+L), F(ab’)2 Fragment (Alexa Fluor® 647 Conjugate; Cell Signaling Technology Cat#4414; 1:500 dilution). After 3 × 10 min washes in 1 × TBS/0.1% Triton X-100 following antibody incubation, the PLVs were mounted on a microscope slide with one drop of both ProLong Gold Antifade Mountant (ThermoFisher Scientific Cat# P36930) and NucBlue Fixed ReadyProbes Reagent (DAPI formulation; ThermoFisher Scientific Cat# R37606). The PLVs were then imaged using a VS120 Slide Scanner for αSMA coverage analysis and Nikon A1R HD confocal microscopy as previously described (*31, 80*).

To validate the mast cell-telocyte interaction, the following primary antibodies targeting mast cell were used in this study: Rabbit anti-human Mast Cell Tryptase IgG (mouse reactivity; Bioss Antibodies Cat# BSM-52533R; 1:100 dilution). To prevent non-specific labeling of Fc receptors (FcR), exclusively secondary antibodies below featuring F(ab) fragments were employed: Anti-rabbit IgG (H+L), F(ab’)2 Fragment (Alexa Fluor® 647 Conjugate; Cell Signaling Technology Cat#4414; 1:500 dilution). After 3 × 10 min washes in 1 × TBS/0.1% Triton X-100 following antibody incubation, the PLVs were mounted on a microscope slide with one drop of both ProLong Gold Antifade Mountant (ThermoFisher Scientific Cat# P36930) and NucBlue Fixed ReadyProbes Reagent (DAPI formulation; ThermoFisher Scientific Cat# R37606). The PLVs were then imaged using a VS120 Slide Scanner for αSMA coverage analysis and Nikon A1R HD confocal microscopy as previously described(*26*).

To confirm telocyte identity and characterize marker expression, whole-mounted lymphatic vessels, OCT-frozen cryosections, or paraffin-embedded tissue sections were subjected to immunofluorescence staining. The following primary antibodies were used: anti-CD34 antibody (Abcam Cat# ab81289; 1:50 dilution), CD31 Monoclonal Antibody (Thermo Fisher Cat# MA1-40074; 1:50), anti-COMP (Abcam Cat# ab231977; 1:125), CD117 (c-Kit) Monoclonal Antibody (Thermo Fisher Cat# 14-1172-81; 1:100), and anti-Myocilin (Abcam Cat# ab41552; 1:50). To prevent non-specific Fc receptor binding, only secondary antibodies with F(ab’) fragments were used: Alexa Fluor® 647–conjugated anti-rabbit IgG (Cell Signaling Technology Cat# 4414; 1:500), Alexa Fluor® 555–conjugated anti-rabbit IgG (Cell Signaling Technology Cat# 4413; 1:1000), FITC-conjugated goat anti-rat IgG (Jackson ImmunoResearch Cat# 112-096-003; 1:200), Alexa Fluor® 647–conjugated donkey anti-rat IgG (Jackson ImmunoResearch Cat# 712-606-153; 1:500), and FITC-conjugated goat anti-rabbit IgG (Jackson ImmunoResearch Cat# 111-096-047; 1:100). Washing, mounting, and imaging were performed as described above.

### Cryosection preparation and Immunohistochemistry (IHC)

Mice were anesthetized with ketamine (100□mg/kg) and xylazine (10□mg/kg) prior to cardiac perfusion with PBS followed by 4% paraformaldehyde (PFA). Tissues of interest were harvested and fixed in 10% neutral buffered formalin (NBF) for 3 days at room temperature. After fixation, tissues were washed three times in PBS and equilibrated in 40% sucrose at 4□°C overnight. Tissues were then infiltrated with optimal control temperature (OCT) embedding medium for 30 minutes at room temperature, transferred to fresh OCT, and snap-frozen for cryosectioning. Frozen blocks were sectioned at 10□μm thickness and collected onto glass slides. Sections were used directly to assess tdT^+^ endogenous expression via fluorescent microscopy or processed for IHC using the same antibodies described above for WHIFM.

### Histochemistry and Histomorphometry

For histochemistry of ankles and knees and quantification of synovitis and osteoclasts, lower limbs of the mice were processed for paraffin embedded demineralized hematoxylin and eosin (H & E) as previously described (*22*). All slides were scanned with an Olympus VS120 (full slide scans available upon request), and histomorphometry for synovial area was quantified with Visiopharm (Hoersholm, Denmark) as previously described (*81*).

### Electron microscopy

Transmission electron microscopy (TEM) was performed as previously described (*20*). The afferent lymphatic vessel to the PLN was identified by injecting Evan’s blue in the footpad and was then excised and fixed overnight at 4°C using a combination fixative of 2.5% glutaraldehyde and 4% paraformaldehyde in 0.1M sodium cacodylate buffer for 24 hours. The specimens were rinsed in 0.1M sodium cacodylate buffer and post-fixed with buffered 1% osmium tetroxide. The tissue was dehydrated in a graded series of ethanol to 100%, transitioned into propylene oxide, infiltrated with EPON/Araldite epoxy resin, followed by embedment in fresh resin and polymerization for 2 days at 70°C. To identify the lymphatic vessel in the specimen, the epoxy embedded block was cut serially into 1μm sections and stained with Toluidine blue. Then the specimen block was trimmed of excess surrounding tissue and thin sectioned at 70 nm with a diamond knife using an ultramicrotome. The thin sections were placed onto 150 mesh carbon coated nickel slot grids and stained with uranyl acetate and lead citrate. A Hitachi 7650 Transmission Electron Microscope with a Gatan 11-megapixel Erlangshen digital camera was used to image the grids. Scanning electron microscopy (SEM) was performed, as previously described.(*26*) Briefly, PLVs were fixed in 2.5% glutaraldehyde/4% paraformaldehyde/0.1M cacodylate overnight, post-fixed in buffered 1% osmium tetroxide, dehydrated, critically point dried, mounted onto aluminum stubs and sputter coated with gold prior to imaging using a Zeiss Auriga FE SEM. Three SEM micrographs per sample group were randomly chosen for descriptive analysis.

### Single-cell RNA sequencing analyses

*In silico* analyses of previously published scRNAseq datasets of WT and TNF-tg PLV (GEO; accession number GSE190999) (*32*) were performed using Seurat (v5.0.1) to generate UMAPs of the mesenchymal cells. Individual gene expression within this UMAP was performed with FeaturePlot in Seurat. Pseudotime predictions analysis of the scRNAseq data from WT and TNF-tg PLV was performed with Monocle 3 to generate UMAPs with heatmaps of the hypothesized cellular differentiation and the telocytes:fibroblasts ratio. For *de novo* scRNAseq of synoviocytes, knee tissue from tamoxifen-treated *Efhd1*CreER x Ai9 mice was digested into a single-cell suspension and processed for scRNAseq as previously described(*32*). RNA-seq libraries were prepared using the NEBNext® Ultra™ II RNA Library Prep Kit plus tdT and sequenced on an Illumina NovaSeq X Plus platform. Reads were aligned to the Mus musculus reference genome (GRCm39) using STAR. UMAPs were generated from the sequence data (GEO; accession number GSE303999) to identify unique cell clusters and expression of marker genes.

### *Ex vivo* co-culture and assays for osmotic pressure response and migration

PLVs with surrounding adipose tissue were harvested from *Myoc*-CreER x Ai9 or *Efhd1*-CreER x Ai9 reporter mice and cultured in DMEM + 10% FBS (Sigma-Aldrich, St. Louis, MO, USA) to establish monolayer co-cultures of tdT^+^ telocytes and unlabeled tdT^−^ fibroblasts. For immunostaining, cells were seeded into the Nunc™ Lab-Tek™ II Chamber Slide™ System (Thermo Fisher Scientific, Cat# 154534PK), fixed with 10% NBF, and the plastic chamber was removed prior to staining. Subsequent staining procedures followed the same protocol as for tissue cryosections. For osmotic shock studies, the primary cells were loaded with Hoechst nuclear stain (Invitrogen, Waltham, MA) and 10 µM Fluo-4 (Invitrogen, Waltham, MA) as previously described (*82*). After 30 minutes of incubation, cells were supplemented with 10 mM Ca^++^ and stimulated with varying concentrations of sucrose (6.25-800 µM) to induce hyperosmotic pressure changes. Real-time fluorescent microscopy was performed to track Ca^++^ influx responses. The percentage of responsive cells was calculated by normalizing the number of Fluo-4^+^ tdT^+^ Hoechst^+^ cells to total tdT^+^ Hoechst^+^ cells for telocytes, and Fluo-4^+^ tdT^−^Hoechst^+^ cells to total tdT^−^ Hoechst^+^ cells for fibroblasts.

### In vitro mt-cpYFP mitoflash assay

Co-cultures of tdTomato-positive (tdT ) telocytes and tdTomato-negative (tdT ) fibroblasts isolated from PLV of tamoxifen-treated *Efhd1*-CreER × Ai9 reporter mice were maintained in DMEM/F-12 medium (Gibco™, Thermo Fisher Scientific, Waltham, MA, USA; Cat# 11320033) supplemented with 10% fetal bovine serum and 1% penicillin–streptomycin. Cells were plated in 35mm Mattek glass bottom dishes (MATTEK, Inc. Ashland, MA, USA; Cat# P35GCOL-1.5-14-C). At approximately 70% confluency, co-cultures were infected with recombinant adenovirus encoding mitochondria-targeted circularly permuted yellow fluorescent protein (Ad-mt-cpYFP (*83*); provided by the laboratory of Dr. Robert T. Dirksen, University of Rochester, Rochester, NY) 48 hours prior to imaging. Immediately before imaging on an inverted Nikon Ti2-E confocal fluorescence microscope, cells were stained with Hoechst to label nuclei and subsequently challenged with 5 mM pyruvate, 100 μM sucrose, or PBS prepared in Krebs-Ringer Solution, HEPES-buffered (Thermo Fisher Scientific, Waltham, MA, USA; Cat# J67795.AP). Time-lapse confocal images were acquired for 3 minutes at a rate of 1.0 s per frame.

### Bulk RNAseq of purified telocyte cultures on stiffness-defined substrates

Telocytes were isolated from the ankle synovium of tamoxifen-treated *Efhd1*CreER x Ai9 mice, and tdT cells were FACS purified and plated onto collagen type I–coated CytoSoft® 24-well plates (Advanced BioMatrix, Cat#5183, #5186, #5189) with defined stiffness levels of 0.2, 8, or 64 kPa. After 24 hours of culture, cells were lysed directly in Buffer RLT Plus (Qiagen, Cat#1053393), and total RNA was extracted using the RNeasy Plus Micro Kit (Qiagen, Cat# 74134) following the manufacturer’s instructions. RNA-seq libraries were prepared using the NEBNext® Ultra™ II RNA Library Prep Kit and sequenced on an Illumina NovaSeq X Plus platform. Reads were aligned to the Mus musculus reference genome (GRCm39) using STAR (GEO; accession number GSE303624). Differential expression analysis was performed using DESeq2, and visualization of gene expression patterns was conducted using the Seurat package (v5.0.1).

### Near infrared-indocyanine (NIR-ICG) imaging and quantification of lymphatic function

The subcutaneous injection of 10 µl of a 0.1 mg/ml ICG solution in water was administered to both hind footpads of the mouse. The fluorescence of the footpad was measured in an IVIS Live Animal Imaging System (Caliper Life Sciences Inc.) one hour after injection (baseline) to capture the maximum fluorescence signal intensity, and again at six hours to quantify the remaining fluorescence signals that were not drained away by lymphatic flow. The ratio of the fluorescence difference and baseline fluorescence was used to define lymphatic clearance function, which reflects the lymphatic vessel’s ability to remove extra fluid from the footpad.

### Zymosan-induced arthritis (ZIA) model

Female 8-weeks-old *Efhd1*-CreER^+⁄–^ x DTA^f⁄–^ mice and wild-type littermate controls received intraperitoneal injections of tamoxifen (0.1 mg/g body weight) for 5 consecutive days to induce *Efhd1*-driven DTA expression and telocyte depletion. Two weeks after the initiation of tamoxifen treatment, mice were anesthetized and received a single intra-articular injection of 180 µg zymosan (Sigma-Aldrich, St. Louis, MO, USA) in 6 µL PBS into the knee joint cavity. Control mice received PBS alone. Knee joint inflammation was monitored by high-resolution ultrasound imaging (Vevo 3100, VisualSonics) performed before zymosan injection (baseline) and on days 2, 7, and 14 post-injection. Analysis of knee joint inflammation was performed using Amira to measure synovial volume, as described before. (*84*) On days 13–15, mice were euthanized, and knee joints were harvested, fixed in 4% paraformaldehyde, micro-CT scanned, and processed for histological analysis.

### Micro-CT analysis

Micro-CT scanning and 3D reconstruction were performed as previously described(*85*), using identical imaging settings and parameters. Following reconstruction, femurs were digitally segmented from adjacent skeletal structures to improve visualization. To assess articular erosions, 2D projections were generated from the articular surface of the femoral trochlea, which articulates with the tibia. This approach maximized the visualization of surface erosion in a flattened plane and minimized distortion-related quantification errors. Erosive lesions were manually annotated using QuPath (v0.5.0), and the cumulative erosion area was normalized to the total projected area of the distal femur (ROI). The erosion percentage (erosion area/ROI) was used as a quantitative measure of joint damage.

### In vitro cellular invasion assays with 3D confocal microscopy and image reconstruction

Matrigel invasion assays were performed using transwell inserts containing cells suspended in serum-free medium. Cells migrated toward a bottom chamber containing 10% fetal bovine serum (FBS), with or without 5 ng/mL TGF-β, for 24 hours. Cultures were then stained with Hoechst to label nuclei and imaged using an inverted Nikon Ti2-E confocal fluorescence microscopy. Representative 3D reconstructions were generated and analyzed using Imaris (version 10.2 Oxford Instruments). For quantitative analysis, the centroid of each Hoechst-stained nucleus was used to define the position of individual cells. Cells were classified as *Efhd1* telocytes if their nuclei were spatially enclosed within the tdTomato signal, or as *Efhd1* fibroblasts if there was no overlap with tdTomato fluorescence. The shallowest nucleus, typically corresponding to a non-migratory or dead cell, was defined as the reference surface level. The vertical distance between each nucleus and this reference plane was calculated as the invasion depth. Distributions of invasion depth were quantified across three independent experiments.

### *Ex vivo* co-culture and assays for osmotic pressure response and migration

PLVs with surrounding adipose tissue were harvested from *Myoc*-CreER x Ai9 or *Efhd1*-CreER x Ai9 reporter mice and cultured in DMEM + 10% FBS (Sigma-Aldrich, St. Louis, MO, USA) to establish monolayer co-cultures of tdT^+^ telocytes and unlabeled tdT^−^ fibroblasts. For immunostaining, cells were seeded into the Nunc™ Lab-Tek™ II Chamber Slide™ System (Thermo Fisher Scientific, Cat# 154534PK), fixed with 10% NBF, and the plastic chamber was removed prior to staining. Subsequent staining procedures followed the same protocol as for tissue cryosections. For osmotic shock studies, the primary cells were loaded with Hoechst nuclear stain (Invitrogen, Waltham, MA) and 10 µM Fluo-4 (Invitrogen, Waltham, MA) as previously described (*82*). After 30 minutes of incubation, cells were supplemented with 10 mM Ca^++^ and stimulated with varying concentrations of sucrose (6.25-800 µM) to induce hyperosmotic pressure changes. Real-time fluorescent microscopy was performed to track Ca^++^ influx responses. The percentage of responsive cells was calculated by normalizing the number of Fluo-4^+^ tdT^+^ Hoechst^+^ cells to total tdT^+^ Hoechst^+^ cells for telocytes, and Fluo-4^+^ tdT^−^Hoechst^+^ cells to total tdT^−^ Hoechst^+^ cells for fibroblasts.

### Statistics

All statistical analyses were performed using GraphPad Prism (version 9.1.1, GraphPad Software, Boston, MA, USA) and R (version 4.3.2). Additional analyses and data visualization, including quantification of osmotic shock assay, invasion assays and ZIA model readouts were conducted using the R packages ggpubr (version 0.6.0). Two-group comparisons were assessed using unpaired two-tailed Student’s t-tests, and multiple group comparisons were performed using one-way or two-way ANOVA as appropriate. A *p*-value less than 0.05 was considered statistically significant.

## Supporting information

Supplementary Video 5 Marked_Telocyte cpYFP 5mM pyruvate002 3min

Supplementary Video 6 Marked_Telocyte mt-cpYFP no stimulate

Supplementary Video 7 Marked_Telocyte cpYFP 100uM sucrose 3min 002

Supplementary Video 2 Antimycin 10min video

Supplementary Video 3 Sucrose 10min video

Supplementary Video 4 NegCtrl 10min video

Supplementary Video 1 Telocyte interacting with mast cells

## Data availability

The data that support the findings of this study are available upon request. Source data are available for Figs. 3c & g, 4b, 5d & f, 6c-f, 9b-c, 10a. The RNA-sequencing datasets have been deposited in GEO database under accession code GSE303999 (scRNA-seq of telocytes in synovial tissue) and GSE303624 (bulk RNA-seq of telocytes cultured under different matrix stiffness conditions).

## Acknowledgements

We would like to thank the faculty and staff at the Genomics Research Center and Histology, Biochemistry, Electron Microscopy Resource in the Center for Advanced Resource Technologies, and Molecular Imaging Core at the University of Rochester Medical Center. This work was supported by funding from the National Institutes of Health: F30 AG069329 (HMK), T32GM007356 (HMK), R01AR063650, R01AG059775 (LX), R01AR069000 (CTR), R01AR056702 (EMS), and P30 AR069655 (EMS).

