## Supplemental Figures for "The Function of *Efhd1*^+^ Telocytes in the Synovial Lymphatic System and Inflammatory-Erosive Arthritis"

| Name | Sequence | Comment |
| --- | --- | --- |
| **Efhd1-CreERT2-F4** | CAA AGG AAG AGG TGT CCT TAG CAG | Genotyping primer for Efhd1CreER strain |
| **Efhd1-CreERT2-R4** | CCT GAA CAT GTC CAT CAG GTT CTT | Genotyping primer for Efhd1CreER strain |
| **Myoc-CreERT2-F4** | CAT CTT GTA CAC GGT GAG CAG | Genotyping primer for MyocCreER strain |
| **Myoc-CreERT2-R4** | GAA GCA TTT TCC AGG TAT GCT CAG AA | Genotyping primer for MyocCreER strain |
| **Pla1a-CreERT2-F4** | CAG CAT GAA GTG CAA GAA CGT G | Genotyping primer for Pla1aCreER strain |
| **Pla1a-CreERT2-R3** | AGA TCA AGG TCT GTA ACA TGA GTC | Genotyping primer for Pla1aCreER strain |

**Supplementary Table 1. Primers used for targeted amplification of fusion genes.**

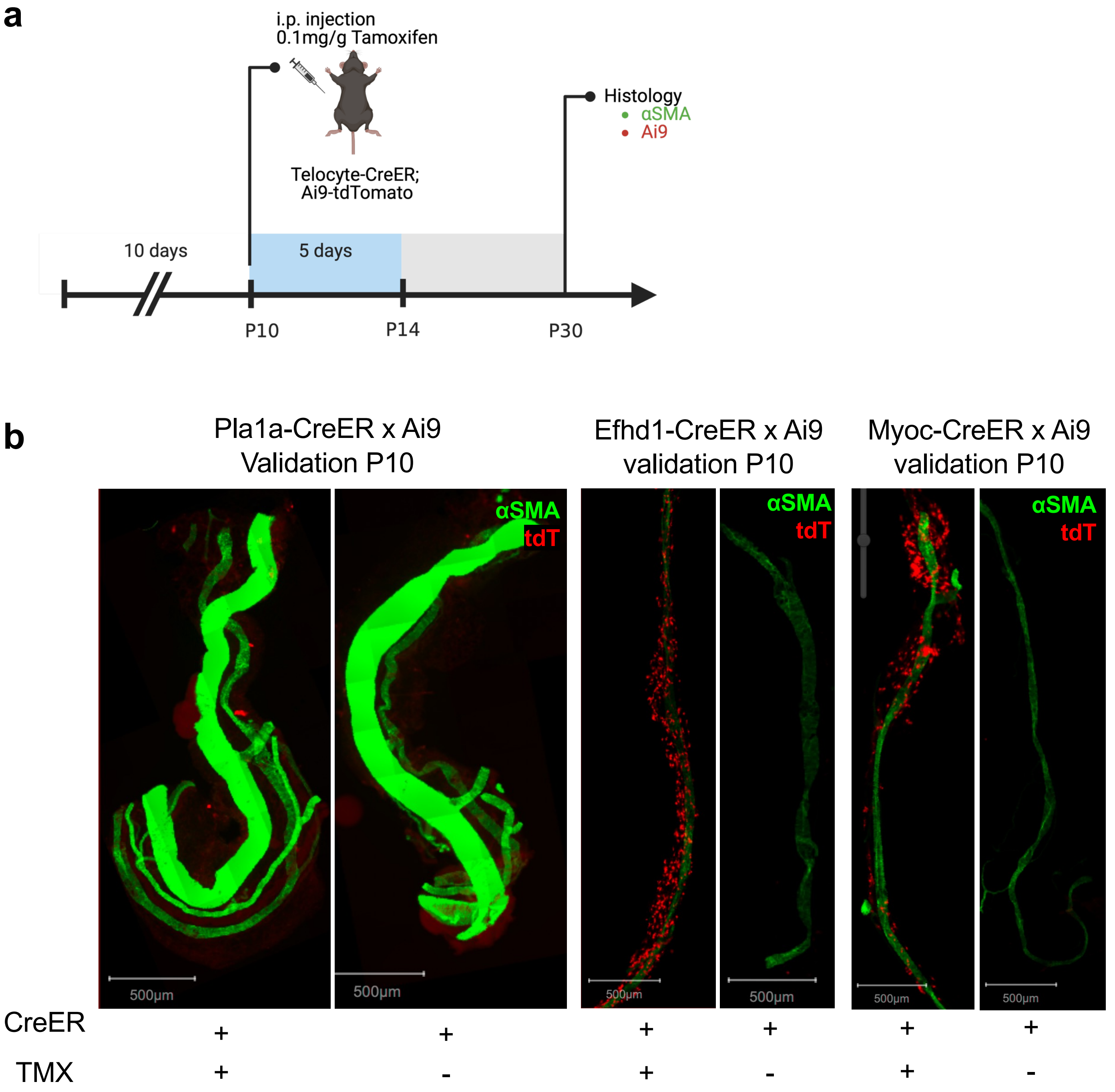


**Supplementary Fig. 1** Validation of Pla1a-CreER, Efhd1-CreER, and Myoc-CreER specificity at early postnatal day 10. **a** Experimental timeline for tamoxifen-induced Cre activation and analysis. Telocyte-CreER;Ai9-tdTomato mice received intraperitoneal injection of tamoxifen (0.1mg/g) at P10, followed by a 5-day chase period until P14, with final dissection and histology analysis at P30. Histological analysis included assessment of αSMA and Ai9 expression. **b** Representative whole-mount images of popliteal lymphatic vessels from Pla1a-CreER x Ai9, Efhd1-CreER x Ai9, and Myoc-CreER x Ai9 mice with and without tamoxifen treatment (TMX). CreER positive status and tamoxifen treatment status are indicated below each image. Data are representative of n ≥3 male and female mice per group.

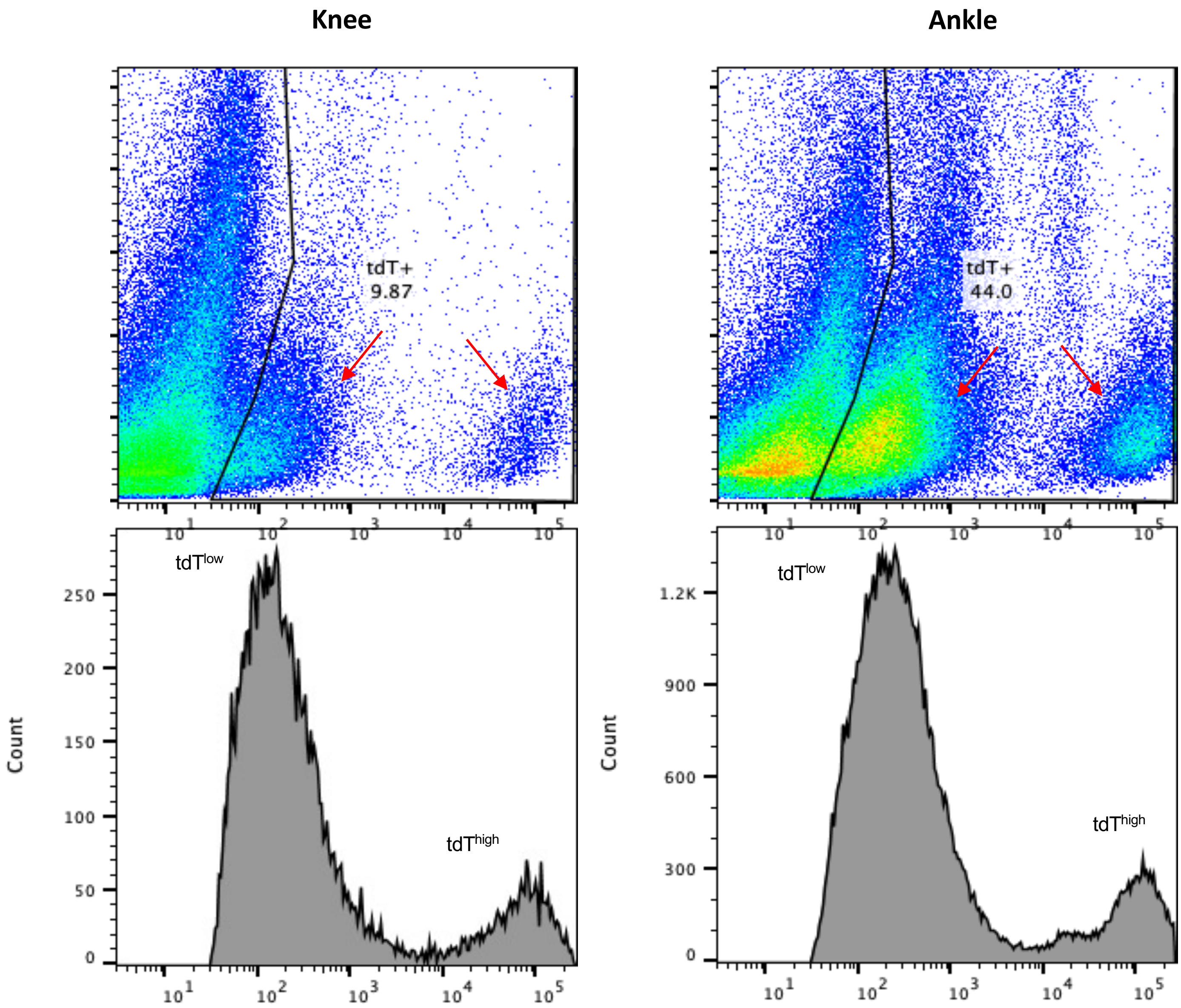


**Supplemental Fig. 2 Flow sorting of tdT$^{+}$ knee and ankle synoviocytes.** Flow cytometric analysis of knee and ankle synovial tissues revealing distinct tdT$^{high}$ and tdT$^{low}$ populations (red arrows). The sorted tdT+ cells from the knee we used for scRNAseq studies.

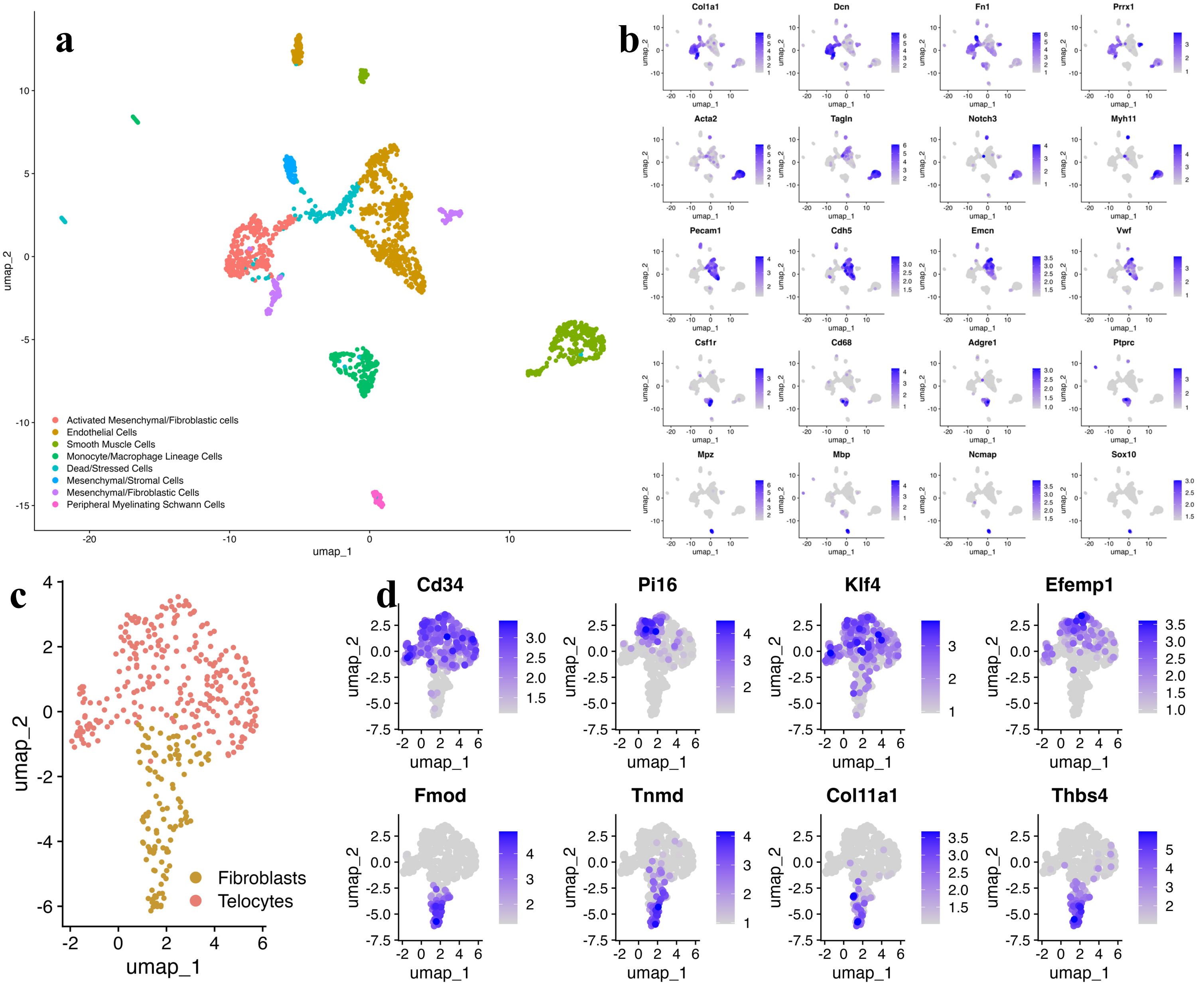


**e**

| Cluster | Marker Gene | Function |
|---|---|---|
| **Telocyte** | Cd34 | Stem/progenitor marker, cell signaling hubs |
| | Pi16 | ECM remodeling and anti-fibrotic functions |
| | Klf4 | Transcription factor, cell plasticity and regulation |
| | Efemp1 | ECM microenvironment modulator and signaling |
| **Fibroblast** | Fmod | Major ECM structural protein, collagen fibrillogenesis |
| | Tnmd | Tendon/ligament development, ECM organization |
| | Col11a1 | Collagen type XI, structural ECM component |
| | Thbs4 | ECM remodeling and cell-matrix interactions |

**Supplemental Fig. 3 a** The scRNAseq data from FACS-purified tdT$^+$ knee synoviocytes described in Fig. 4b were used to generate a UMAP where a minimally supervised shared nearest neighbor clustering algorithm resolved 13 clusters, defined by marker gene expression. For downstream analysis, only clusters corresponding to Mesenchymal/Fibroblastic cells and Mesenchymal/Stromal cells were retained, while clusters such as Smooth Muscle Cells, Endothelial Cells, Monocyte/Macrophage Lineage Cells, and Peripheral Myelinating Schwann Cells were excluded. **b** UMAPs showing the relative expression levels of representative marker genes confirm the identity of each cluster. **c** A focused UMAP of the retained Mesenchymal/Fibroblastic and Mesenchymal/Stromal cells is presented, highlighting two dominant populations distinguished by their specialized roles in microenvironmental regulation (Telocytes) and extracellular matrix production (Fibroblasts). **d** Expression plots display the levels of genes involved in extracellular matrix production and microenvironmental regulation across these populations. **e** Telocyte and Fibroblasts clusters are defined by functional genomics.

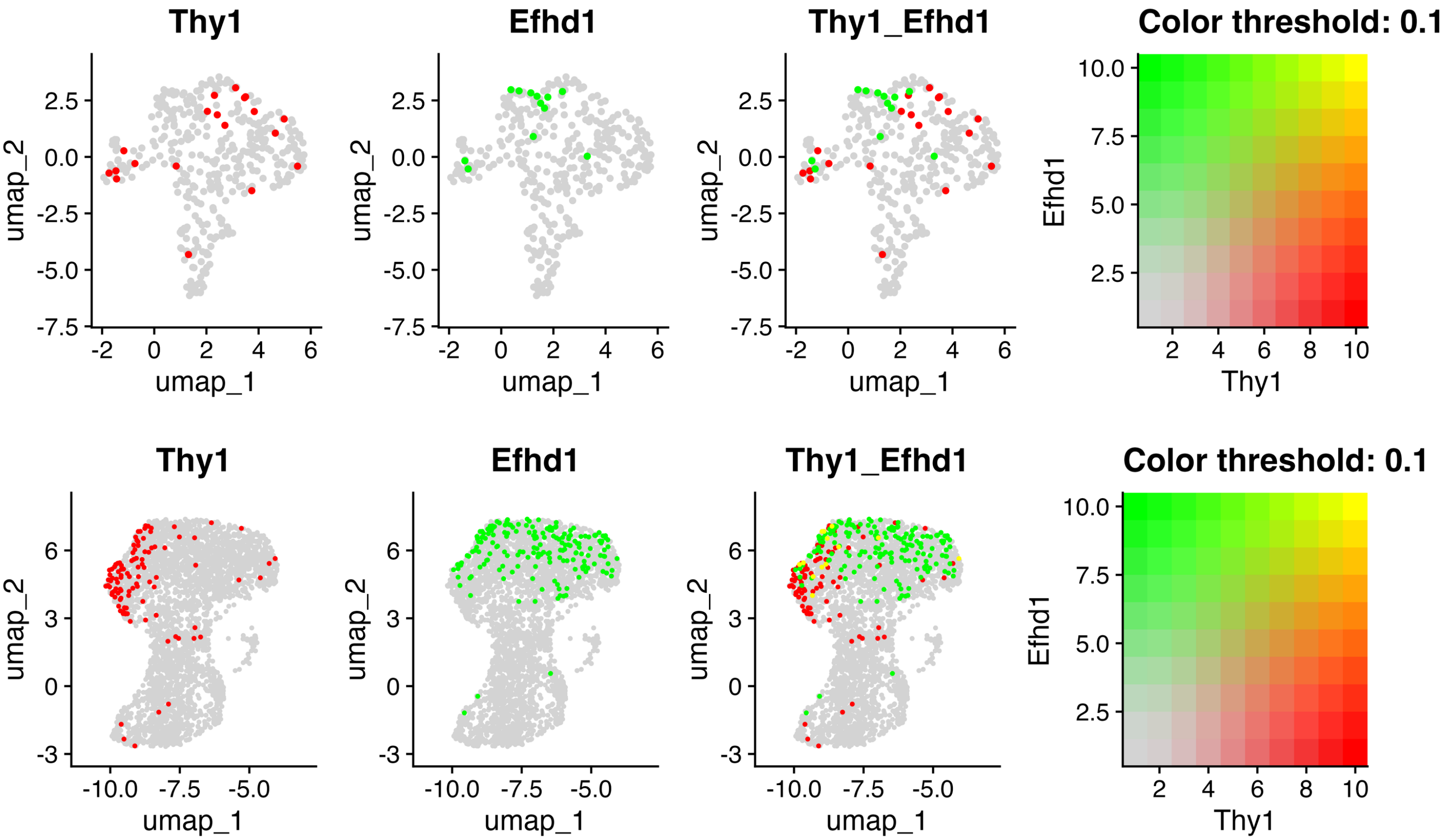


**Supplemental Fig. 4 Non-overlapping expression of *Thy1* and *Efhd1* in knee synovium tissues and PLV.** Single cell expression of *Thy1* and *Efhd1* superimposed on the UMAPs described in Figure 3 and 4.

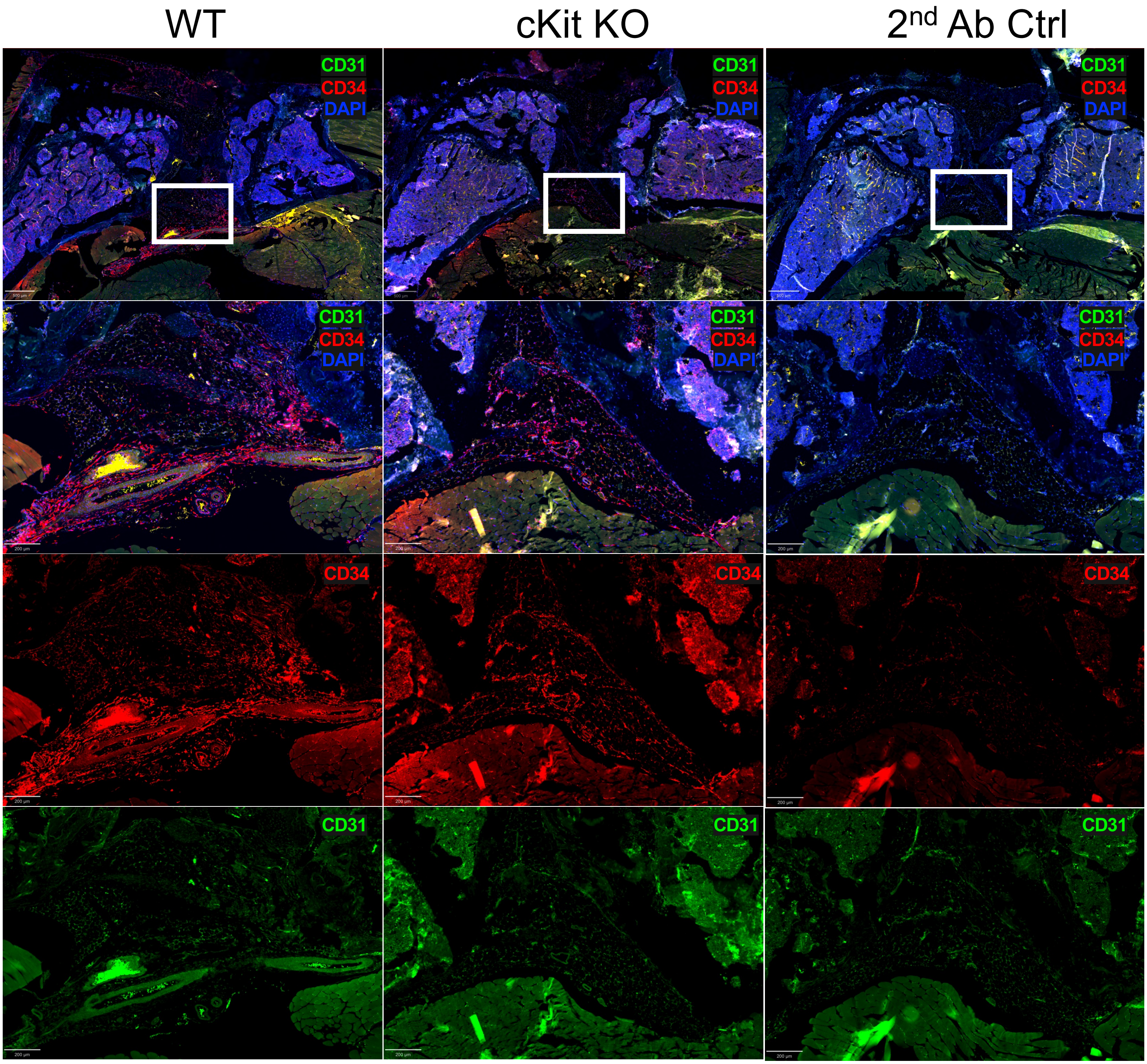


**Supplemental Fig. 5 Identification of CD34$^{+}$/CD31$^{-}$ telocytes in the knee synovium of cKit$^{-/-}$ mice.** IHC with labelled antibodies and DAPI counter stain was performed on the knee histology sections from WT and cKit$^{-/-}$ mice, and representative fluorescent microscopy images are shown highlighting CD31 (green) and CD34 (red) expression, with DAPI counter stain (blue). Top row: 4x magnification images reveal similar numbers of CD34$^{+}$/CD31$^{-}$ telocyte-like cells within the synovial tissues of WT and cKit$^{-/-}$ mice. Bottom row: 10x magnification views of boxed regions confirm the identity of CD34$^{+}$/CD31$^{-}$ synovial telocytes.

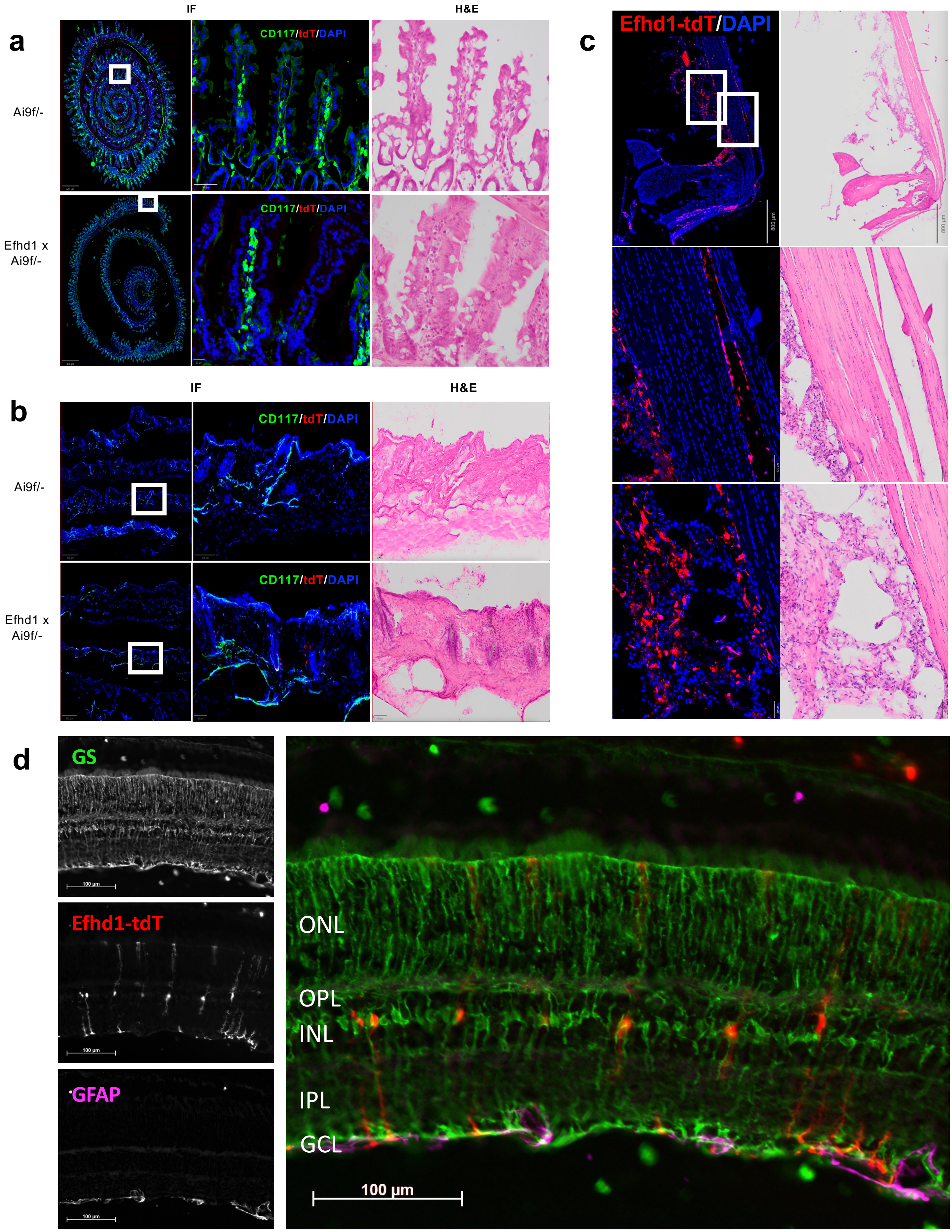

a
IF
H&E
Ai9f/-
Efhd1 x Ai9f/-
CD117/tdT/DAPI
CD117/tdT/DAPI
b
IF
H&E
Ai9f/-
Efhd1 x Ai9f/-
CD117/tdT/DAPI
CD117/tdT/DAPI
c
Efhd1-tdT/DAPI
d
GS
Efhd1-tdT
GFAP
100 µm
ONL
OPL
INL
IPL
GCL
100 µm

**Supplemental Fig. 6  Efhd1-CreER$^{+/-}$x Ai9$^{+/-}$ mice mediate tamoxifen-induced gene expression in tissue-specific telocytes.** Fresh-frozen **a** intestine, **b** skin, and **c** Achilles tendon tissues were harvested from tamoxifen-treated Efhd1-CreER$^{+/-}$x Ai9$^{+/-}$ mice and processed for DAPI counter-stained IHC with labelled antibodies against CD117/cKit for fluorescent microscopy, and subsequent H&E stained for brightfield microscopy.  Representative 4x and 10x images are shown to illustrate: the abscess of tdT$^{+}$ (red) cells in intestine including CD117+ ICC, tdT$^{+}$ telocytes in hair follicles, and tdT$^{+}$ telocytes at synovial-tendon attachments.  **d** Fresh frozen retina sections were processed for IHC with labelled antibodies against glutamine synthetase (GS; green) and glial fibrillary acidic protein (GFAP; magenta). Note the small percentage of GS+ Müller glia are also tdT$^{+}$.  In contrast, no tdT+ telocytes in the intestine were detected. Similar results were obtained with tissues from Myoc-CreER$^{+/-}$x Ai9$^{+/-}$ mice (data not shown).

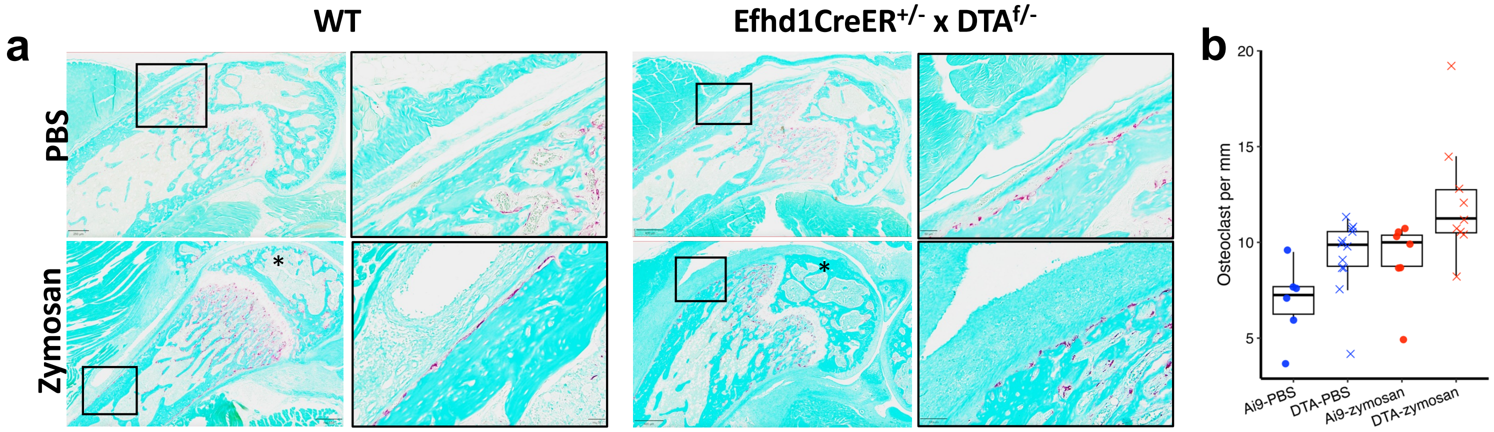


**Supplemental Fig. 7 Telocyte depletion effects on cortical bone osteoclasts and focal erosions from zymosan induced arthritis (ZIA)**. **a** Representative TRAP-stained histology slides obtained at 10x from the experiment described in Figure 6 are presented with magnified ROI to highlight TRAP$^{+}$ osteoclasts (red) on the femoral cortical bone surface. Histomorphometry was performed to quantify these osteoclasts as described in Methods. Of note is that no osteoclasts were observed in the histology (*) corresponding to the patella grove focal erosions observed in the micro-CT images, confirming that this component of the acute ZIA response was completed by day 14 post-injection. Similar results were obtained in experiments with Myoc-CreERT2$^{+/-}$ x DTA$^{f/-}$ mice (data not shown).

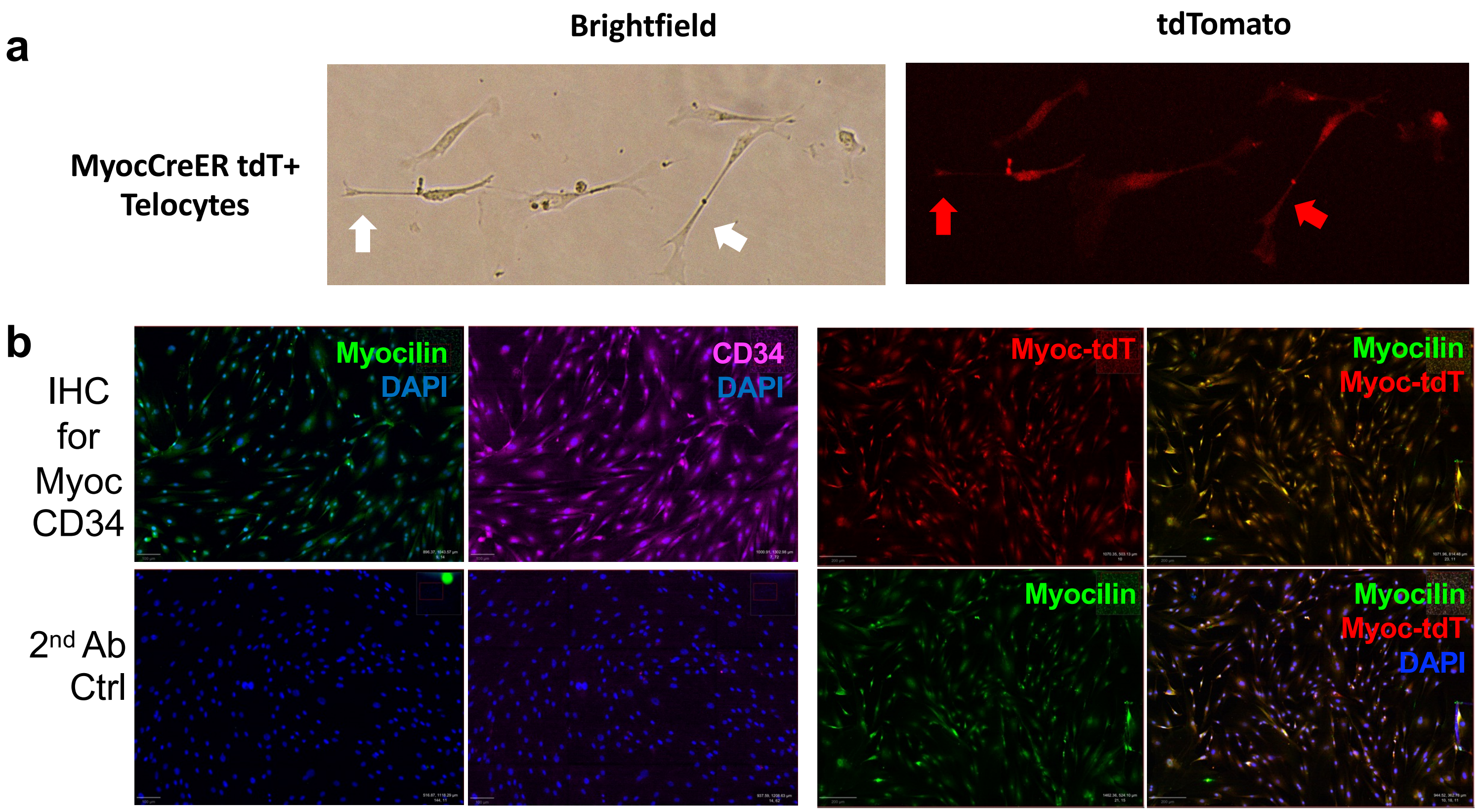


**Supplementary Fig. 8 Validation of FACS purified telocytes in cultures.** FACS purified $tdT^+$ cells described in Fig. 4 were cultured in complete growth DMEM medium supplement with F12. **a** Representative brightfield with fluorescent 20x micrographs of 4th passage cells in culture are shown highlighting characteristic telopodes (arrows). **b** Cultured $tdT^+$ telocytes were seeded in chamber slides and subjected to immunofluorescent staining with labelled antibodies against Myocilin (green) and CD34 (magenta), and counter stained with DAPI (blue) prior to fluorescent microscopy. Representative 5x images are shown (top panels) with $2^{nd}$ antibody controls (bottom panels). Note colocalization of the Myocilin immunoreactivity with tdT (yellow cells) confirming that the culture cells retain their telocyte phenotype.

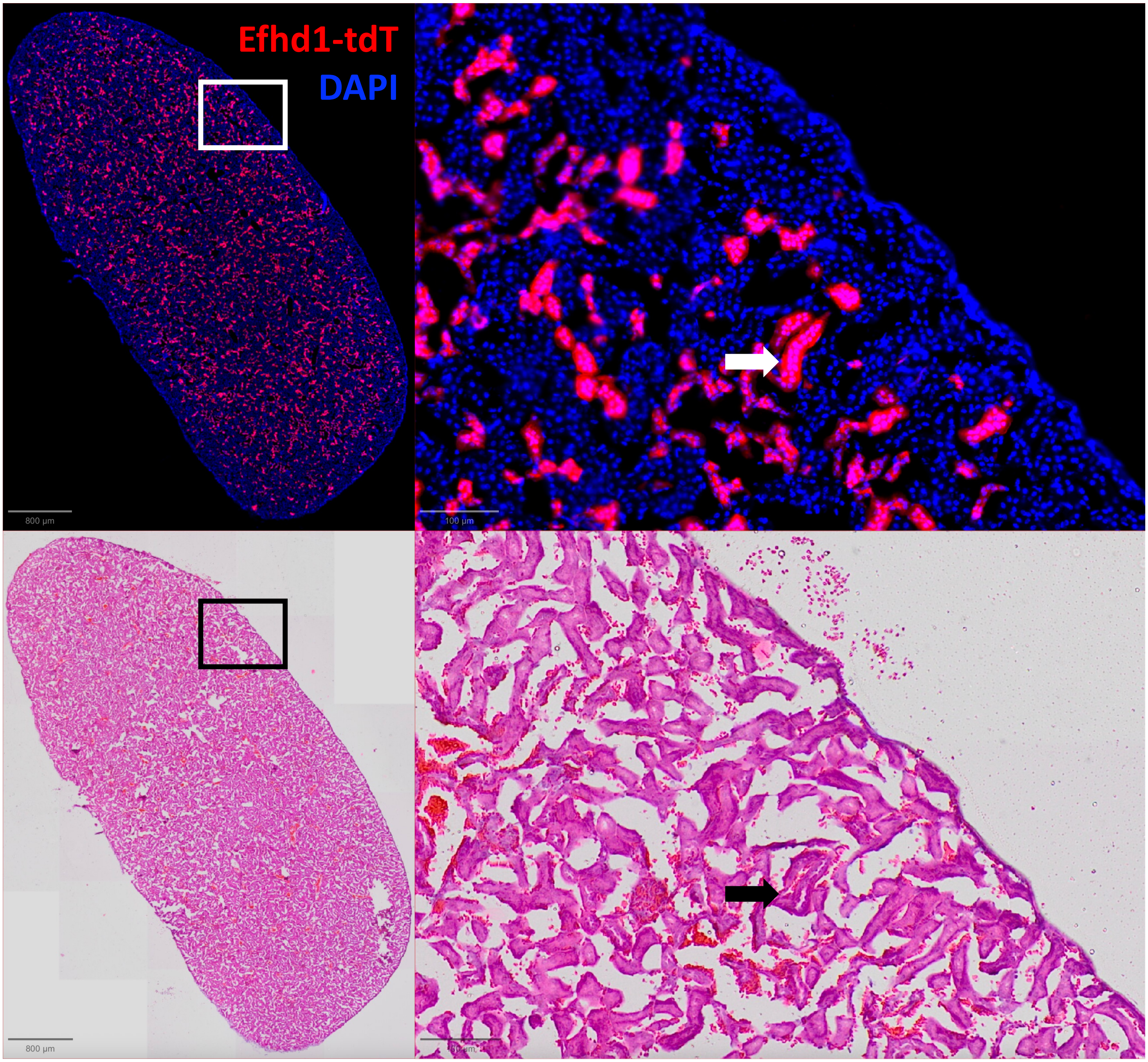


**Supplemental Fig. 9 Efhd1-CreER targeting of renal tubules in kidney.** Fresh-frozen histology sections of kidney tissue from tamoxifen treated Efhd1-CreER$^{+/-}$x Ai9$^{+/-}$ mice were processed and counterstained with DAPI for fluorescent microscopy, and representative 4x image with 10x image of the boxed region of interest are shown (top panels). The slides were then H&E stained for brightfield microscopy (bottom panels). Arrow indicate the tdT$^{+}$ renal tubules.
